# Development of Comprehensive Ultraperformance Liquid Chromatography-High Resolution Mass Spectrometry Assays to Quantitate Cisplatin-Induced DNA-DNA Cross-Links

**DOI:** 10.1101/2022.09.28.509855

**Authors:** Arnold S. Groehler, Asema Maratova, Nhat Mai Dao, Anuar Mahkmut, Orlando D. Schärer

**Author notes:** Corresponding author (Orlando D. Schärer).

## Abstract

Cisplatin (CP) is a common anti-tumor drug used to treat many solid tumors. The activity of CP is attributed to the formation of DNA-DNA cross-links, which consist of 1,2-intra-, 1,3-intra-, and interstrand cross-links. To better understand how each intrastrand cross-link contributes to the activity of CP, we have developed comprehensive ultraperformance liquid chromatography-selective ion monitoring (UPLC-SIM) assays to quantify 1,2-GG, 1,2-AG, 1,3-GCG, and 1,3-GTG-intrastrand cross-links. The limit of quantitation for the developed assays ranged from 5 – 50 fmol, or as low as 6 cross-links per 10^8^ nucleotides. To demonstrate the utility of the UPLC-SIM assays, we first performed *in vitro* cross-link formation kinetics experiments. We confirmed 1,2-GG-intrastrand cross-links were the most abundant intrastrand cross-link and formed at a faster rate compared to 1,2-AG- and 1,3-intrastrand cross-links. Furthermore, we investigated the repair kinetics of intrastrand cross-links in CP-treated wild type and nucleotide excision repair (NER)-deficient U2OS cells. We observed slow repair of both 1,2- and 1,3-intrastrand cross-links in wild type cells, and no evidence of repair in the NER-deficient cells. Taken together, we have demonstrated that our assay is capable of accurately quantifying intrastrand cross-links in CP-treated samples and can be utilized to better understand the activity of CP.

## INTRODUCTION

The anti-tumor drug *cis*-diamminedichloroplatinum(II) (cisplatin, CP) is a common chemotherapeutic agent used as a first-line treatment against many solid tumors such as breast, ovarian, and testicular cancers^1-3^. The anti-tumor activity of cisplatin is generally attributed to the formation of DNA-DNA cross-links. If left unrepaired, these cross-links can interfere with crucial biological processes such as DNA replication and transcription, ultimately leading to apoptosis and cell death. Despite its clinical utility, cisplatin treatment can yield serious side effects including gastrointestinal and renal toxicity, neuropathy, and ototoxicity^1^. Furthermore, the development of cisplatin-resistance has been reported in patients across different cancers^4^.

Cisplatin is comprised of a platinum atom bound to two amine groups and two chloride groups susceptible to displacement. After entering the cell, the difference in chloride ion concentration in the cellular matrix compared to the cytoplasm stimulates the nonenzymatic displacement of chloride groups with water^2^. This activated species can react with nucleophilic positions on DNA yielding a monoadduct, which can then form intra- or interstrand cross-links with adjacent bases (**Scheme 1**)^5^. The most abundant cisplatin-induced cross-link is the 1,2 guanine-guanine intrastrand cross-link (CP-d(GG)), comprising an estimated 65 – 75% of all DNA adducts^6–8^. The remaining cisplatin-induced cross-links are attributed to 1,2 adenine-guanine intrastrand cross-links (CP-d(AG), estimated 15 – 25%), 1,3 guanine-X-guanine intrastrand cross-links (CP-d(GXG), where X can be 2’-deoxycytidine or thymidine, estimated 5 – 10%), and interstrand cross-links (ICLs, estimated <1%)^6–8^. The formation of cisplatin-induced DNA-protein cross-links have been identified as a minor product as well^6, 9–10^.

**Scheme 1:**
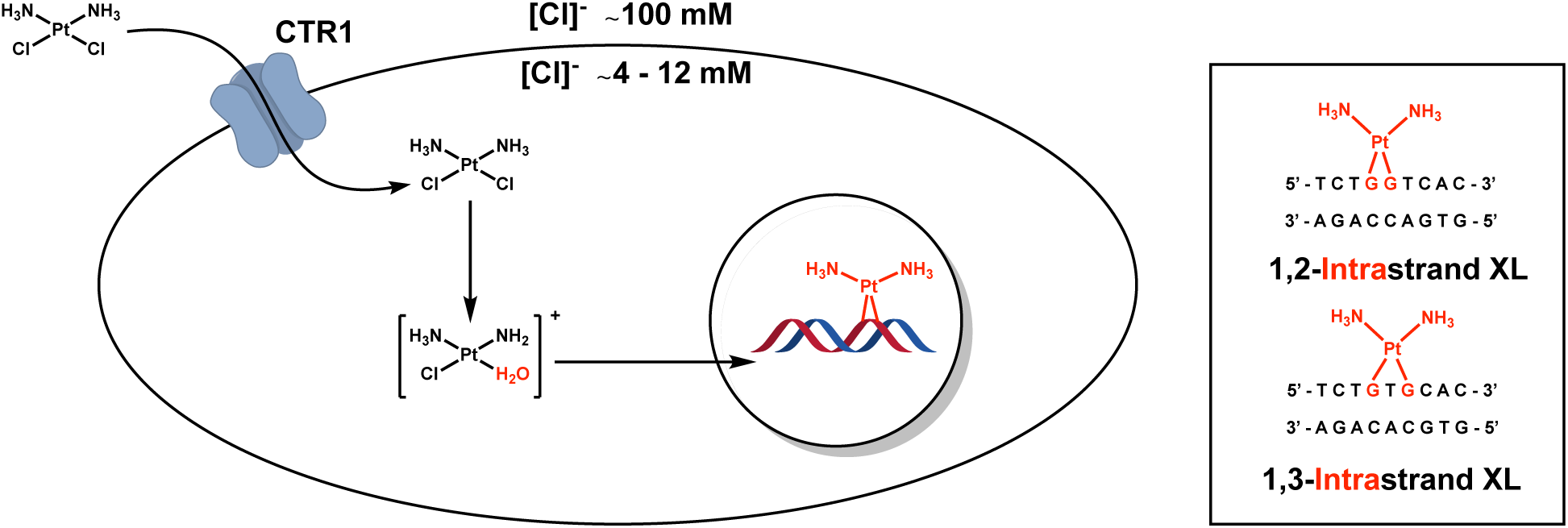
Cisplatin mechanism of action

Given that DNA adducts have the potential to be excellent biomarkers for monitoring cisplatin dosing *in vivo*, influx/efflux of drug into cells, and the formation/repair of DNA damage, many researchers have attempted to develop quantitative assays to monitor platinum-induced DNA-DNA crosslinking. Previously developed assays have relied upon atomic absorption spectroscopy (AAS)^11^, antibody probes against Pt-DNA damage^12^, and ^32^P post-labeling^13^ of Pt-nucleosides following DNA digestion. AAS can accurately detect and quantitate platinum levels but is not sensitive enough to measure complex biological samples. Immunoblotting assays are attractive due to their simplicity and applicability to most laboratories, but the currently available CP-d(GG) antibodies are prone to cross-reactivity and thus yield non-linear responses^12^. Radiolabeling CP-d(XG) and CP-d(GXG) with ^32^P is sensitive enough to detect as adduct levels as low as 0.087 fmol DNA adduct per µg DNA (2.7 adducts per 10^8^ nucleotides), but requires the use of radioactive material and the technique is labor intensive^13^. Importantly, all three methodologies fail to differentiate the structure of the DNA adduct, making it impossible to correlate interstrand, 1,2- and 1,3-intrastrand crosslink levels to the biological response. The distinction of DNA adducts is of particular importance because the levels of repair of interstrand and intrastrand adducts, by homologous recombination/BRCA-dependent^14–16^ and nucleotide excision repair-dependent pathways^17–19^, respectively, have been implicated in influencing the success of therapeutic outcomes.

More recently, inductively coupled plasma mass spectrometry (ICP-MS) and liquid chromatography tandem mass spectrometry (HPLC-MS/MS) assays have been developed to accurately quantitate individual CP-induced adducts. Garcia *et al* developed a HPLC-ICP-MS assay to quantitate the most abundant CP-d(GG) in *Drosphilia melanogaster* somatic cells^20^. However, their assay had a reported limit of quantitation (LOQ) of 1 CP-d(GG) per 10^6^ nucleotides and was unable to measure CP-d(GG) at cisplatin treatments below 500 µM^20^. Alternatively, Baskerville-Abraham *et al* developed a HPLC-MS/MS assay with a LOQ of 3.7 CP-d(GG) per 10^8^ nucleotides, capable of quantitating CP-d(GG) from 12.5 µM CP-treated cells and tissue samples from mice treated with 7 mg/kg cisplatin^21^. Unfortunately, neither methodology was able to accurately detect or quantitate 1,2-AG intrastrand cross-links or 1,3-intrastrand cross-links. Finally, Henderson *et al* developed accelerator mass spectrometry assays to quantify cross-linking by ^14^C-labeled platinum drugs carboplatin^22^ and oxaliplatin^23–24^. Although accelerator mass spectrometry affords superior sensitivity (1 ± 0.1 amol cross-link per µg DNA), and allows analysis at sub-phamacological levels, their methodologies once again failed to differentiate different 1,3-intrastrand cross-links from ICLs. Since 1,3-intrastrand cross-links are better substrates for nucleotide excision repair, quantitation of 1,3-intrastrand cross-links is better suited for investigating the role of DNA repair in cisplatin toxicity/resistance. Furthermore, comparison of both 1,2- and 1,3-intrastrand cross-links holds more potential to accurately correlate DNA adduct formation and persistence to the mechanism of action of cisplatin.

To address this need, we have developed a comprehensive ultraperformance liquid chromatography-high resolution mass spectrometry-selective ion monitoring (UPLC-SIM) assay to quantitate 1,2- and 1,3-intrastrand cross-links in cisplatin-treated calf thymus DNA and cell culture samples. We were able to accurately quantitate the repair of cisplatin-induced intrastrand cross-links in cisplatin-treated nucleotide excision repair (NER)-deficient and isogenic control cells, validating our methodology for more detailed studies of the repair of cisplatin adducts. Our developed assay has the potential to accurately quantitate CP-d(GpX) in cells and tissue biopsies, with the view of informing cisplatin regiments in the context of personalized medicine.

## MATERIALS AND METHODS

### Caution

*Cisplatin is a known carcinogen and should be handled with appropriate personal protective equipment (ie. gloves and laboratory coats) should be worn. All generated waste should be disposed following environmental regulations*.

### Chemicals

Unless otherwise stated, all chemicals and enzymes were purchased from Merck & Co. (Kenilworth, NJ). Isotopically labeled ^15^N_5_-labeled phosphoramidites were purchased from Cambridge Isotope Laboratories (Tewksbury, MA). All reagents used for DNA oligonucleotide synthesis were purchased from Glen Research (Sterling, VA). Exonuclease I (Exo I), exonuclease III (Exo III), exonuclease V (Exo V), T7 exonuclease (T7 Exo), Quick calf intestinal alkaline phosphosphatase (Quick CIP), MNase, and nucleoside digestion mix were purchased from New England Biolab (NEB, Ipswich, MA). DNase was purchased from Roche (Basel, Switzerland). Nanosep 10KDa centrifugal filters were purchased from Pall Corporation (New York, NY). Ultra-pure (UP) water was purchased from Biosesang (Seongnam, South Korea).

### Preparation of CP-d(GpX) analyte standards

Authentic standards of CP-d(GpX) digestion products were synthesized for UPLC-SIM assay development and optimization. 5’-3’ GG, AG, GA, GCG, and GTG dimers and trimers were prepared on a 1µM scale using standard DNA oligo synthesis protocols (5’-trityl protecting group removed) on a Mermade 4 automated DNA synthesizer and deprotected with 55% NH_4_OH. HPLC purification was performed using an Agilent 1260 Infinity HPLC coupled with an Agilent 1260 Infinity II photodiode array detector and a Phenomenex Clarity 5 μm Oligo-RP (150 x 4.6 mm) column. A gradient of 100mM triethylamine acetate (TEAA, buffer A) and 100% methanol (buffer B) was operated at 1 mL/min starting at 2% B for 2 min, linearly increased to 9% B over 10 minutes, then 25% B over 10 minutes, then 80% B over 6 minutes, held constant at 80% B for 2 minutes, followed by a decrease to 2% B over 1 minute, and finally re-equilibrated at 2% B for 9 minutes. Under these conditions, AG dimer, GA dimer, GG dimer, GCG trimer, and GTG trimer eluted at 22.1, 21.2, 19.3, 21.4, and 22.8 min respectively.

HPLC-purified dimers and trimers were characterized by liquid chromatography-mass spectrometry (LC-MS) with a full-MS negative mode assay using a Q-Exactive Focus mass spectrometer coupled to a Dionex Ultimate 3000 UPLC system as follows; A Thermo Hypersil-Gold 1.9 μm C18 column (100 x 2.1 mm) was operated using a gradient of 15mM ammonium acetate, pH 7.0 (buffer A) and 100% Acetonitrile (buffer B) at 0.05 mL/min starting at 2% B for 2 min, linearly increased to 80% B over 16 minutes, held constant at 80% B for 2 minutes, followed by a decrease to 2% B over 2 minutes, and finally re-equilibrated at 2% B for 12 minutes. MS settings were as follows; Scan range 150 – 2000 *m/z*, electrospray voltage (3000 V), automatic gain control (AGC, 1e^6^), capillary temperature 320 °C, HESI temperature 150 °C, sheath gas, auxiliary gas, and sweep gas flow rate 35, 10, and 1 arbitrary units respectively. ESI^-^-MS (GG): *m/z* (−1) = 595.1400; ESI^-^-MS (AG or GA): *m/z* (−1) = 579.1444; ESI^-^-MS (GCG): *m/z* (−1) = 884.1856; ESI^-^-MS (GTG): *m/z* (−1) = 889.1853. Stock concentrations of each dimer and trimer were determined using UV absorbance measured on a microvolume UV spectrophotometer (Thermo Scientific^™^ NanoDrop 2000/2000c) using the following extinction coefficients: AG ε_260_ = 25000, GA ε_260_ = 25200, GG ε_260_ = 21600, GCG ε_260_ = 28200, and GTG ε_260_ = 30300.

Cisplatin was activated by incubating 2.4 mg (0.008 mMol) cisplatin and 1.3 mg (0.0076 mMol) silver nitrate in 1 mL UP-water overnight at 37 °C protected from light. After incubation, the resulting silver chloride precipitate was removed using a 0.2 μm nylon filter yielding an 8mM solution of activated cisplatin. In an Eppendorf tube, dimer or trimer oligonucleotides were incubated with 1 equivalent of activated cisplatin overnight at 37 °C protected from light. CP-d(GpX) was purified by the HPLC method described above with CP-d(GA), CP-d(GG), CP-d(AG), CP-d(GCG), and CP-d(GTG) eluting at 11.5, 14.0, 17.0 (broad), 17.3, and 19.6 minutes respectively.

Each CP-d(GpX) standard was characterized by a UPLC-parallel reaction monitoring (UPLC-PRM) assay in positive mode as follows; A Waters HSS T3 1.9 μm C18 column (100 x 2.1 mm) was operated using a gradient of 15mM ammonium acetate, pH 7.0 (buffer A) and 100% methanol (buffer B) at 0.05 mL/min starting at 2% B for 2 min, linearly increased to 25% B over 8 minutes, followed by an increase to 50% B over 20 minutes, then an increase to 80% over 2 minutes, held constant at 80% for 2 minutes, followed by a decrease to 2% B over 2 minute, and finally re-equilibrated at 2% B for 15 minutes. MS settings were as follows; Electrospray voltage (3000 V), capillary temperature (320°C), full scan AGC (1e^6^), full scan resolution 70,000, HESI temperature 150°C, sheath gas, auxiliary gas, and sweep gas flow rate 35, 10, and 1 arbitrary units respectively, PRM AGC (5e^4^) and PRM resolution 35,000. Due to the natural occurrence of platinum isotopes, the three most abundant *m/z* were analyzed by UPLC-PRM as follows. ESI^+^-PRM CP-d(GG): *m/z* (+2) = 412.08214, 412.58366, and 413.08330 from 15 – 17.5 min; ESI^+^-PRM CP-d(GA) and CP-d(AG): *m/z* (+2) = 404.08554, 404.5868, and 405.08699 from 14.5 – 16.5 or 17.5 – 19.5 min respectively; ESI^+^-PRM CP-d(GCG): *m/z* (+2) = 556.60529, 557.10616, and 557.60675 from 16 – 18 min; ESI^+^-PRM CP-d(GTG): *m/z* (+2) = 564.1100, 564.6060, and 565.10643 from 17.5 – 19.5 min.

### Preparation of ^15^N_5_-labeled CP-d(GpX) internal standards

Isotopically labeled internal standards of ^15^N_5_-CP-d(GG), ^15^N_5_-CP-d(AG), ^15^N_5_-CP-d(GCG), and ^15^N_5_-CP-d(GTG) were synthesized and characterized analogously as CP-d(GpX) standards. In short, ^15^N_5_-labeled 5’-3’ GG, AG, GCG, and GTG dimers and trimers were prepared using a Mermade 4 automated DNA synthesizer. The coupling of the ^15^N_5_-labeled nucleotide was performed manually by adding 200 μL 67 mM 2’-deoxyadenosine (^15^N_5_) or 2’-deoxyguanosine (^15^N_5_) phosphoramidite as the 5’-terminal guanine or adenine. Dimer/trimer HPLC purification, CP-d(GpX) synthesis, and purification was performed exactly as described above for the authentic standards.

Each ^15^N_5_-CP-d(GpX) internal standard was characterized by the UPLC-PRM assay in positive mode described above for the authentic standards with the following changes; ESI^+^-PRM ^15^N_5_-CP-d(GG): *m/z* (+2) = 414.57290, 415.07618, and 415.57572 from 15 – 17.5 min; ESI^+^-PRM ^15^N_5_-CP-d(AG): *m/z* (+2) = 406.57441, 407.07910, and 407.57941 from 17.5 – 19.5 min; ESI^+^-PRM ^15^N_5_-CP-d(GCG): *m/z* (+2) = 559.09391, 559.59858, and 560.09917 from 16 – 18 min; ESI^+^-PRM ^15^N_5_-CP-d(GTG): *m/z* (+2) = 566.59730, 567.09842, and 567.59885 from 17.5 – 19.5 min.

### Synthesis of 42mer intrastrand cross-link substrate

To a labeled eppendorf tube, 44.8 nmol of 42mer oligo was diluted to 1345 μL 10 mM sodium perchlorate (NaClO_4_) and 5 mM acetic acid and treated with one equivalent of aquated cisplatin for 1 hour at 37°C protected from light. The sequence of the oligos were as follows; 5’-TCT TCT TCT TCT TCT TCT **GG**T TCT TCT TCT TCT TCT TCT TCT-3’ (42mer-GG XL), 5’-TCT TCT TCT TCT TCT TCT **AG**T TCT TCT TCT TCT TCT TCT TCT-3’ (42mer-AG XL), 5’-TCT TCT TCT TCT TCT TCT **GCG** TCT TCT TCT TCT TCT TCT TCT-3’ (42mer-GCG XL), and 5’-TCT TCT TCT TCT TCT TCT **GTG** TCT TCT TCT TCT TCT TCT TCT-3’ (42mer-GTG XL).

After incubation, precipitated cisplatin was removed using a 0.2 μm NYLON filter. The 42mer 1,2-intra or 1,3-intrastrand cross-link oligo substrate was purified using an AKTA Pure fast protein liquid chromatography (FPLC) as follows; A MonoQ 5/50 GL column was operated using a gradient of (A) 10mM NaOH and (B) 1M NaCl in 10 mM NaOH at 2 mL/min starting at 10% B for 10 column volumes (CV) to equilibrate, then kept at 10% B for 5 CV after sample injection, followed by increasing to 30% B over 5 CV, then to 50% B over 40 CV, then to 100% B over 20 CV, held constant for an additional 5 CV, and then decreased to 10% B to re-equilibrate for 10 CV. Under these conditions, the 42mer 1,2- and 1,3-intrastrand cross-link substrate eluted between 61 – 64 CV, while the unplatinated 42mer oligo eluted later between 64 – 66 CV. All collected substrates were concentrated to dryness using lyophilization.

Once dried, the resulting solids were resuspended in 1 mL of UP-water and desalted using centrifugal filtration (Merck, Amicon^®^ Ultra 10 kDa filters) at 14,000 rcf for 10 minutes at 4 °C. The samples were washed with UP-water an additional three times to ensure buffer exchange. The products were confirmed by the negative mode full-scan LC-MS method described above.

### Evaluation of digestion enzymes

A 100 pmol aliquot of 42mer oligo containing a site-specific 1,2- or 1,3-intrastrand cross-link was incubated in the presence of a single exonuclease. Digestion conditions, enzyme concentrations, and digestion times were optimized during analysis. The digestion reactions were quenched by adding an equal volume of bromophenol blue in 90% formamide and then resolved on a 20% urea gel with 1X TBE buffer at 300 volts for 2.5 hours. The resulting digestion products were visualized by staining with SYBR Gold and analyzed with an Amersham^™^ Typhoon^™^ biomolecular imager (GE Healthcare Life Sciences).

As additional confirmation of digestion efficiency, the 42mer oligo digestions above were repeated and the reaction products were HPLC purified using the CP-d(GpX) method described above. Under these conditions, the desired 24mer or 21mer containing a cisplatin 1,2- or 1,3-intrastrand cross-link eluted as a broad peak between 25 – 28 min.

### Platination of calf thymus DNA (CTDNA) with activated cisplatin

A stock solution of CTDNA was resuspended in water at a concentration of 1 mg/mL. CTDNA (500 μg) was reacted with an increasing range of aquated cisplatin (50 nm, 100 nm, 250 nm, 500 nm, 1 μM, 2 μM, 5 μM, and 10 μM, N=3), brought to a final volume of 650 μL, and incubated at 37 °C for 24 hours protected from light. After incubation, excess cisplatin was removed by centrifugal filtration (Merck, Amicon^®^ Ultra 3 kDa filters) at 14,000 rcf for 10 minutes at 4 °C. The platinated CTDNA was further washed with an equal volume of UP-water three additional times, followed by recovering the CTDNA following the manufacturer’s protocol. Platinated CTDNA concentration was measured using the double-strand DNA (dsDNA) settings on a microvolume UV spectrophotometer (Thermo Scientific^™^ NanoDrop 2000). Purified platinated CTDNA solutions were stored at -20 °C.

### Cross-link formation kinetics in cisplatin-treated CTDNA

A stock solution of 5 mM cisplatin was prepared in 0.9% NaCl and vortexed for 30 minutes to ensure everything was dissolved immediately before the experiment. CTDNA (250 μg) was reacted with the cisplatin solution above to yield a final concentration of 500 nM (final volume 1250 μL) and incubated at 37 °C for 1, 2, 4, 6, 24, or 48 hours (N=4) protected from light. After incubation, three replicates were quenched by incubating with 10 mM thiourea (final volume 1500 μL) at 37 °C for an additional 1 hour. After quenching, excess cisplatin and thiourea was removed by centrifugal filtration (Merck, Amicon^®^ Ultra 3 kDa filters) at 14,000 rcf for 10 minutes at 4 °C. The platinated CTDNA was further washed with an equal volume of UP-water three additional times, followed by precipitating the DNA with a 2X volume of 100% EtOH. The precipitated CTDNA was pelleted by centrifugation at 20,000 rcf for 10 minutes at 4 °C, then washed with 70% EtOH 2Xs and 100% EtOH 1X with the centrifugation conditions above. The remaining replicate was treated exactly as described above without the thiourea quenching. Platinated CTDNA concentration was measured using the double-strand DNA (dsDNA) settings on a microvolume UV spectrophotometer (Thermo Scientific^™^ NanoDrop 2000). Purified platinated CTDNA solutions were stored at -20 °C.

### Digestion and sample enrichment of CTDNA for *in vitro* quantitation of CP-d(GpX)

Aliquots of platinated CTDNA (25 μg to 100 μg) were spiked with 1 pmol of CP-d(GpX) IS, followed by enzymatically digesting the samples to CP-d(GpX) analytes using the following procedures. While optimizing digestion methods, technical replicates (N=2 or 3) were analyzed to confirm reproducibility. After digestion optimization for each CP-d(GpX), CTDNA platination described above was repeated to obtain experimental replicates (N=3) of CP-d(GpX) quantitation.

*Method 1*: A 50 μg aliquot of CP-treated CTDNA was diluted to 320 μL of 50 mM sodium acetate and 10 mM MgCl_2_ and treated with 5 U of DNaseI for 4 hours at 37 °C. After 4 hours, 5 U of Nuclease P_1_ (NucP_1_) was added, and the solution was incubated an additional 16 hours at 37 °C. Finally, 41 μL 1M Tris-HCl, pH 9.0 and 5 U of Quick CIP (NEB, Ipswich, MA) was added, and the solution was incubated an additional 4 hours at 37 °C, followed by enzyme removal and sample enrichment as described below.

*Method 2*: A 50 μg aliquot of CP-treated CTDNA was diluted to 150 μL of 50 mM sodium acetate and 10 mM MgCl_2_ and treated with 5 U of DNaseI for 4 hours at 37 °C. The solution was diluted to 200 μL to yield 1X NEBuffer 1 (NEB, Ipswich, MA) at pH 7.0, and 50 U of Exo III was incubated 4 hours at 37 °C, followed by adding 0.05 U PDE II and incubating 16 hours at 37 °C. Finally, 41 μL 1M Tris-HCl, pH 9.0 and 5 U of Quick CIP (NEB, Ipswich, MA) was added, and the solution was incubated for an additional 4 hours at 37 °C, followed by enzyme removal and sample enrichment as described below.

*Method 3*: A 25 μg aliquot of CP-treated CTDNA was diluted to 100 μL of 1X nucleoside digestion mix reaction buffer (NEB, Ipswich, MA) at pH 5.4 and treated with 2.5 μL NEB nucleoside digestion mix (1 μL or 1 U per 10 μg CTDNA) for 10 minutes at 37 °C. Immediately after digestion, the reaction was quenched by addition of 10 mM EDTA, followed by enzyme removal and sample enrichment as described below.

*Method 4*: A 50 μg aliquot of CP-treated CTDNA was diluted to 100 μL of 4 mM MgCl_2_ and treated with 25 U of benzonase and 1.75 U Quick CIP for 24 hours at 37 °C. After 24 hours, the solution was diluted to 150 μL to yield 1X Nuclease S_1_ (NucS_1_) buffer (Thermo, Waltham, MA), and 10 U of NucS_1_ was incubated an additional 1 hour at 37 °C. Finally, enzymes were removed and the samples enriched as described below.

*Enzyme removal*: The digestion enzymes of each reaction were removed by centrifugation at 14,000 rcf for 10 minutes at 4 °C through Nanosep^®^ centrifugal devices with Omega^TM^ 10kDa membranes. The filters were further washed using an equal volume of DI water one additional time, and 100 μL 50:50 ACN:UP-water two additional times. The combined elution fractions were combined and concentrated to dryness by centrifugal vacuum.

*HPLC enrichment*: CP-d(GpX) were enriched by HPLC purification using an Agilent 1260 Infinity HPLC coupled with an Agilent 1260 Infinity II photodiode array detector and a Phenomenex Clarity 5 μm Oligo-RP (150 x 4.6 mm) column following the methodology described above for CP-d(GpX) authentic standards. During analysis, fractions were collected in 1-minute intervals from 11.0 – 22.0 minutes using the automated fraction collector. Fractions from 11.0 – 14.0 minutes (CP-d(GG)), 16.0 – 19.0 minutes (CP-d(AG) and CP-d(GCG)), and 19 – 21 minutes (CP-d(GTG)) were pooled together and concentrated by vacuum centrifugation and lyophilization. The concentrated samples were resuspended in 10 μL LC-MS water for UPLC-SIM analysis.

### Ultra-performance liquid chromatography-single ion monitoring (UPLC-SIM) CP-d(GpX) assays

Each CP-d(GpX) analyte was analyzed by a developed UPLC-SIM assay in positive mode as follows; A Waters HSS T3 1.9 μm C18 column (100 x 1.0 mm) was operated using a gradient of 15mM ammonium acetate, pH 7.0 (buffer A) and 100% methanol (buffer B) at 0.05 mL/min starting at 2% B for 2 min, linearly increased to 25% B over 8 minutes, followed by an increase to 50% B over 10 minutes, then an increase to 80% over 2 minutes, held constant at 80% for 2 minutes, followed by a decrease to 2% B over 2 minute, and finally re-equilibrated at 2% B for 9 minutes. MS settings were as follows; Electrospray voltage (3000 V), capillary temperature 320 °C, HESI temperature 150 °C, Sheath gas, auxiliary gas, and sweep gas flow rate 35, 10, and 1 arbitrary units respectively.

Due to the natural occurrence of platinum isotopes, the two most abundant *m/z* were detected for both the analyte and internal standard, respectively, by UPLC-SIM as follows. ESI^+^-SIM CP-d(GG): *m/z* (+2) = 412.58366, 413.08330, 415.07618, and 415.57572 from 9.0 – 13.0 min; ESI^+^-SIM CP-d(AG): *m/z* (+2) = 404.5868, 405.08699, 407.07910, and 407.57941 from 11.5 – 16.5 or 11.0 – 15.0 min respectively; ESI^+^-SIM CP-d(GCG): *m/z* (+2) = 557.10616, 557.60675, 559.59858, and 560.09917 from 11.0 – 15 min; ESI^+^-SIM CP-d(GTG): *m/z* (+2) = 564.6060, 565.1064, 567.0984, and 567.5989 from 12.0 – 16.0 min.

To avoid any potential, theoretical overlap of analyte and internal standard isotope signals, the third isotope signal (^196^Pt) was used for quantitation and the more abundant second isotope signal (^195^Pt) was used for confirmation (**Table 1**). For low abundance samples from cisplatin-treated cells, the second isotope signal was used for quantitation. CP-d(GpX) quantitation was performed by dividing the area under the analyte signal by the internal standard signal and multiplying by the amount of internal standard spiked into the sample.

**Table 1:** Representative CP-d(GpX) analyte and IS *m/z* isotope distributions Observed *m/z* (+2) of each authentic standard and internal standard used for confirmation and quantitation.

| Structure | Analyte confirmation | Analyte quantitation | IS confirmation | IS quantitation |
| --- | --- | --- | --- | --- |
| CP-d(GG) | 412.5836 | 413.0833 | 415.0762 | 415.5757 |
| CP-d(AG) | 404.5868 | 405.0870 | 407.0791 | 407.5794 |
| CP-d(GCG) | 557.1062 | 557.6067 | 559.5986 | 560.0992 |
| CP-d(GTG) | 564.6060 | 565.1064 | 567.0984 | 567.5989 |

### CP-d(GpX) UPLC-SIM assay validation

Aliquots of 25 µg CTDNA were spiked with 1 pmol CP-d(GG), CP-d(AG), CP-d(GCG), or CP-d(GTG) IS and the following amounts of authentic standards (N = 3): 500, 250, 100, 40, or 20 fmol CP-d(GG); 1000, 500, 400, 250, or 100 fmol CP-d(AG); 500, 200, 100, 50, or 20 fmol CP-d(GCG) or CP-d(GTG). For 1,2-intrastrand cross-links and 1,3-intrastrand cross-links, CTNDA was digested by *Method 1* and *Method 3,* respectively, exactly as described above, followed by processing and enrichment by Nanosep 10kDa filtration and HPLC purification. Collected fractions were analyzed by UPLC-SIM as described above, and the observed ratio of analyte to internal standard was plotted against the expected ratios. Accuracy of each point was calculated by subtracting the mean ratio by the expected ratio, followed by dividing by the expected ratio. The precision of each point was calculated by dividing the standard deviation by the mean ratio. The limit of detection of each assay was determined as a signal to noise ratio of 3:1, and the limit of quantitation was determined as a signal to noise ratio of 10:1 from the validation results. Interday validation was performed by repeating the experiment above, and by analyzing the individual sample sets on separate days.

### Cisplatin treatment of cell culture

U2OS Flp-In/T-REX cells WT and with XPA knocked out CRISPR/Cas9^25^ were seeded (1.0 x 10^6^ cells) overnight in Dulbecco’s Modified Eagle’s Medium (DMEM) supplemented with 10% fetal bovine serum and 1% penicillin-streptomycin at 37 °C and 5% CO_2_. Cells were cultured and allowed to grow to 70% confluency prior to treatment.

A stock solution of 5.0 mM Cisplatin was prepared in 0.9% sodium chloride solution and vortexed 30 minutes to ensure everything was dissolved. The cell media was removed, cells were washed twice with phosphate-buffered saline, and cell media with 10 µM or 25 µM cisplatin was added. After 2 hours of incubation, the media was again removed, and cells were washed two times with PBS. Fresh cell media was added, and the cells were incubated an additional 1, 2, 4, 8, or 24 hours to allow repair of CP-induced DNA cross-linking. After the appropriate repair time, cells were trypsinized and collected for future processing. For the “zero-hour repair” sample, cells were harvested immediately after the two-hour incubation with cisplatin.

DNA was extracted by first lysing collected cells in Qiagen Cell Lysis Solution (1 mL per 1.0 x 10^7^ cells). Each solution was supplemented with 70 U RNaseA and incubated at 37 °C for 5 hours. After RNA digestion, protein was digested by adding 8 U of proteinase K and incubating at 37 °C for an additional 1 hour. The proteinase K was precipitated by adding 200 µL Qiagen Protein Precipitation Solution and vortexing for 30 seconds. The precipitated proteins were pelleted by centrifugation at 5000 rpm for 10 minutes at 4 °C. The supernatant was decanted to labeled 15 mL tornado tubes and genomic DNA was precipitated by adding two volumes of 100% ethanol. Precipitated DNA was pelleted by centrifugation at 3900 RPM for 30 minutes at 4 °C. The pelleted DNA was washed, digested, and processed using the appropriate methodology described above.

## RESULTS

### Workflow for method development for the analysis of CP-d(GpX) adducts in calf thymus and cellular DNA

Our approach to the detection and quantification is based on the digestion of DNA containing the various cisplatin DNA intrastrand crosslinks into nucleotide dimers and trimers of distinct structure and molecular weight, which can be first separated by HPLC and detected by UPLC-SIM. Thus, we anticipated that our method would be able to distinguish and allow quantification of the four main cisplatin intrastrand adducts: CP-d(GG), CP-d(AG), CP-d(GCG), and CP-d(GTG). This required (i) the generation of isotopically labeled internal standards for quantification, (ii) development of a method a UPLC-SIM method for their detection, and (iii) method for the efficient digestion into the nucleotide dimers and trimers while avoiding the over digestion of the internal phosphodiester bonds.

### Synthesis and characterization of CP-d(GpX) standards and internal standards

Authentic standards and ^15^N_5_-labeled internal standards of CP-d(GG), CP-d(AG), CP-d(GCG), and CP-d(GTG) were synthesized by reaction of the appropriate dimer or trimer oligonucleotide with activated cisplatin (**Figure 1**). Each CP-d(GpX) was HPLC purified and characterized by UV, high resolution accurate mass-mass spectrometry (HRAM-MS), and UPLC-PRM. The isotope purity of the ^15^N_5_-labeled internal standards were confirmed using UPLC-SIM as described in materials and methods.

**Figure 1:**
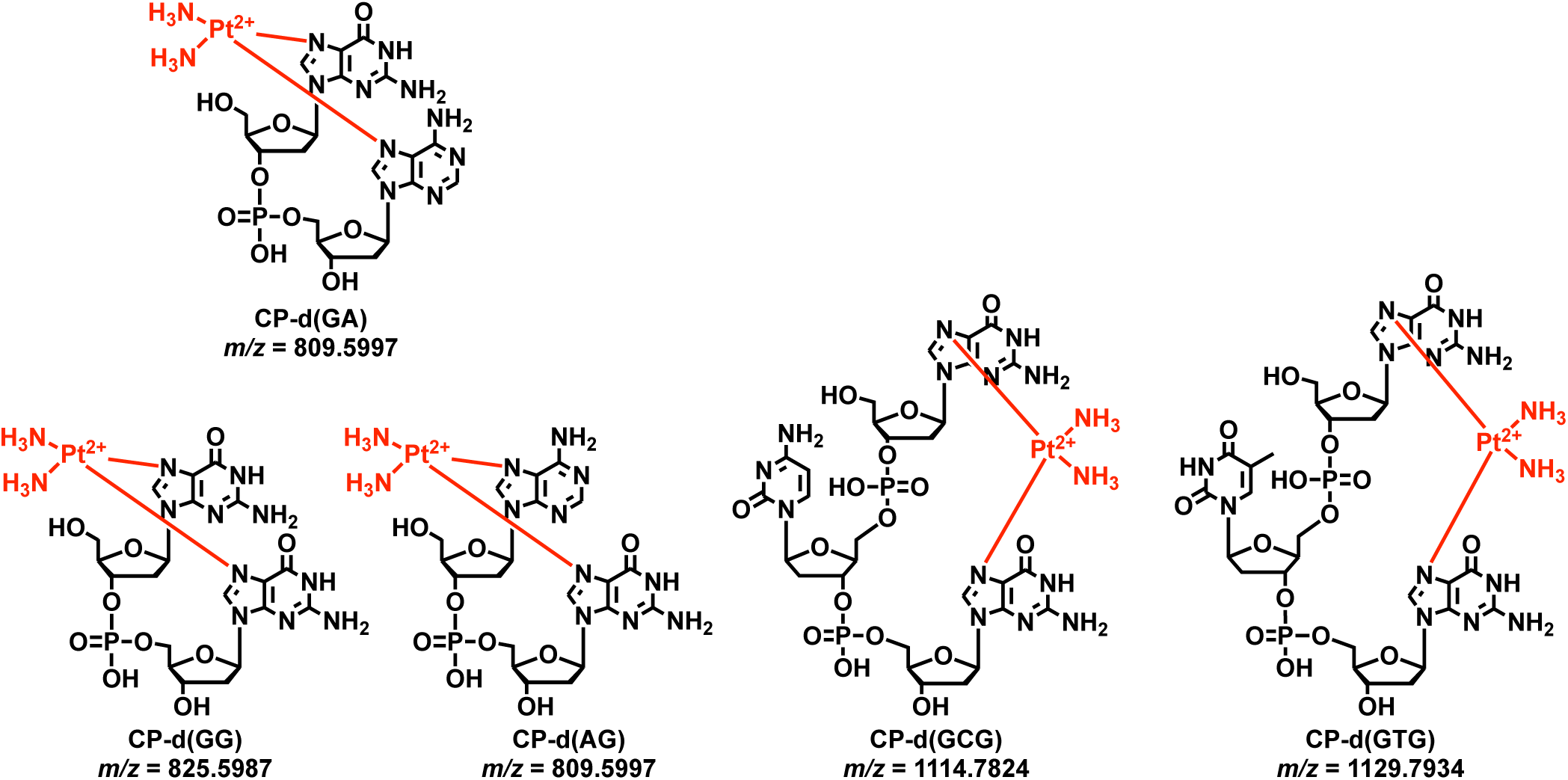
Chemical structures of CP-d(GG), CP-d(AG), CP-d(GA), CP-d(GCG), and CP-d(GTG) analytes derived from digestion of DNA containing 1,2-intrastrand and 1,3-intrastrand cross-links

Each CP-d(GpX) authentic standard and internal standard was initially analyzed by UPLC-HRAM-MS in negative and positive modes because each species can exist as a zwitterion, with the negative charge on the phosphate group(s) and the positive charge on the N7 position of adenine or guanine. This analysis revealed much greater sensitivity in the positive mode monitoring +2 ionization states for all CP-d(GpX) investigated. Furthermore, the UPLC-HRAM-MS confirmed the expected platinum isotope distribution of at least 6 isotopes (three major and three minor, **Figure 2A**, **Figure S1**), which was utilized in the UPLC-SIM development to provide additional confidence in product identification.

**Figure 2:**
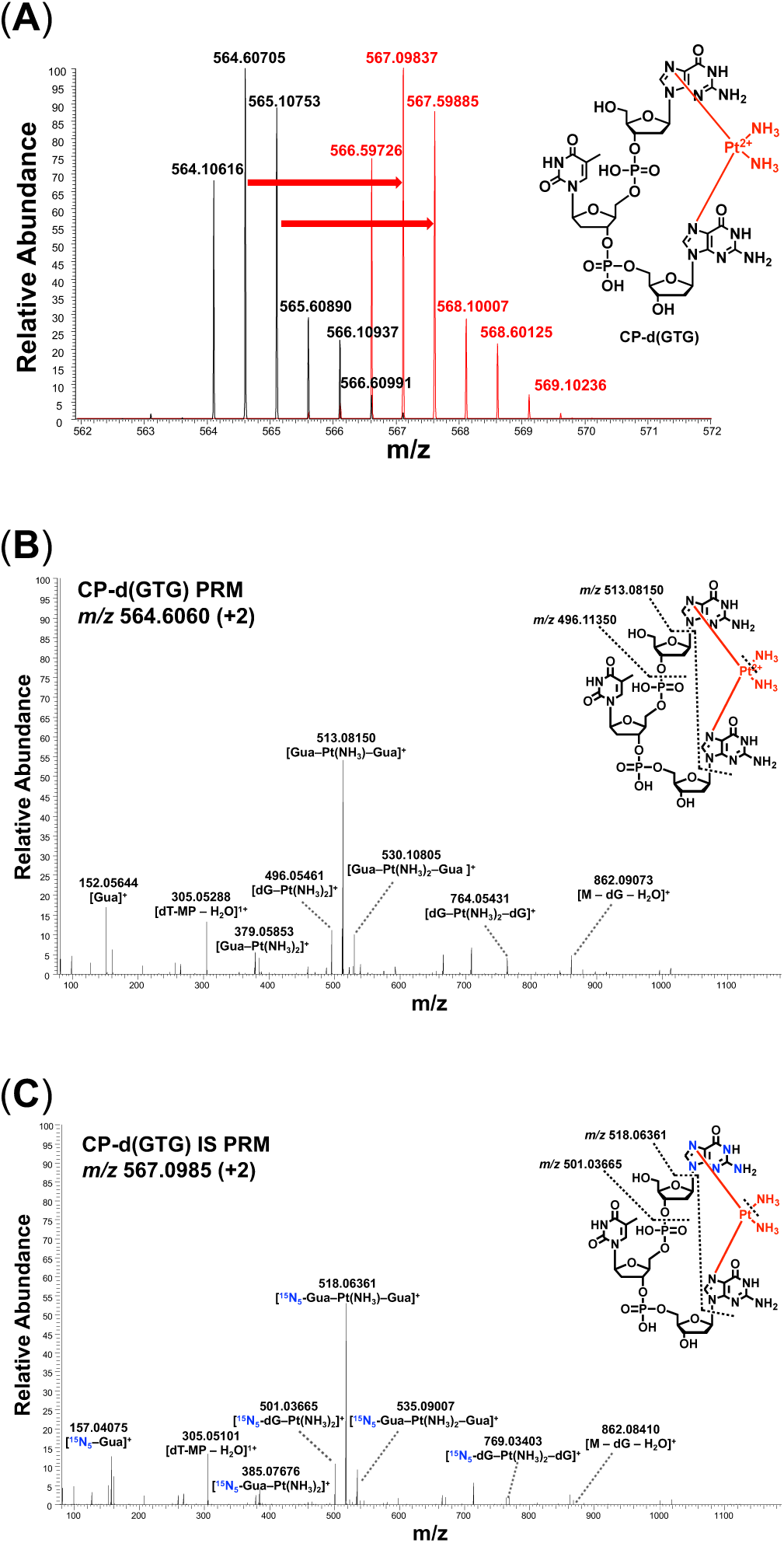
Characterization of CP-d(GpX) standards and internal standards. (**A**) Representative isotope distribution pattern of CP-d(GTG) analyte (black) and ^15^N_5_-CP-d(GTG) internal standard (red). (**B**) Representative fragmentation pattern of the CP-d(GTG) analyte. (**C**) Representative fragmentation pattern of the ^15^N_5_-labeled CP-d(GTG) internal standard. The ^15^N_5_-isotopes are highlighted in blue.

CP-d(GpX) were further characterized by UPLC-PRM fragmentation on the three most abundant isotopes for each authentic standard (**Figure 2B** & **S2**, **Table S1**) and internal standard (**Figure 2C** & **S3**, **Table S2**). For CP-d(GG), CP-d(GCG), and CP-d(GTG) the major fragments were breakage of the glycosidic bonds and one or two amine groups to yield two guanines cross-linked by platinum ([Gua–Pt(NH_3_)_x_–Gua]^+^ where X = 0, 1, or 2). For example, the most abundant isotope of the CP-d(GTG) analyte yielded fragments of *m/z* (+1) = 496.0543, *m/z* (+1) = 513.0813, and *m/z* (+1) = 530.1079 corresponding to breakage of the glycosidic bonds and loss of two, one, and zero amine groups respectively (**Figure 2B**). Additional minor fragments included the M/2 state of the major fragments above, 5’-guanine cross-linked to platinum ([Gua–Pt(NH_3_)_2_], *m/z* (+1) = 379.0585), and two 2’-deoxyguanosines cross-linked to platinum ([dG–Pt(NH_3_)_2_–dG], *m/z* (+1) = 764.0543). UPLC-PRM analysis of the ^15^N_5_-labeled internal standards yielded analogous fragments with the expected *m/z* shift (**Figure 2C**). UPLC-PRM of CP-d(GA) and CP-d(AG) also induced breakage of the platinum bond to the guanosine, yielding a single nucleoside [Ade-Pt(NH_3_)_2_]^+^ as a major fragment (**Figure S2**).

### UPLC-SIM assay development

Using the synthesized and characterized authentic and ^15^N_5_-labeled internal standards, we investigated different UPLC-HRAM-MS conditions. All CP-d(GpX) are extremely polar and did not retain well on traditional C8 or C18 columns. The Waters HSS T3 (150 x 1.0 mm, 1.9 µm) column was the only column investigated that provided reasonable retention profiles of CP-d(GG), CP-d(AG), CP-d(GCG), and CP-d(GTG) eluting at 10.6, 13.1, 12.3, and 13.6 minutes respectively (**Figure 3A**). To increase overall sensitivity, we investigated the effect of lowering the pH of the 15 mM ammonium acetate buffer from 7.0 to 6.0 and 4.5 to increase the percentage of doubly charged species. Using stock solutions of CP-d(GpX) standard/IS, we observed a dramatic decrease in sensitivity with lowering the pH of the buffer, confirming pH 7.0 was the optimal condition (**Table S3**).

**Figure 3:**
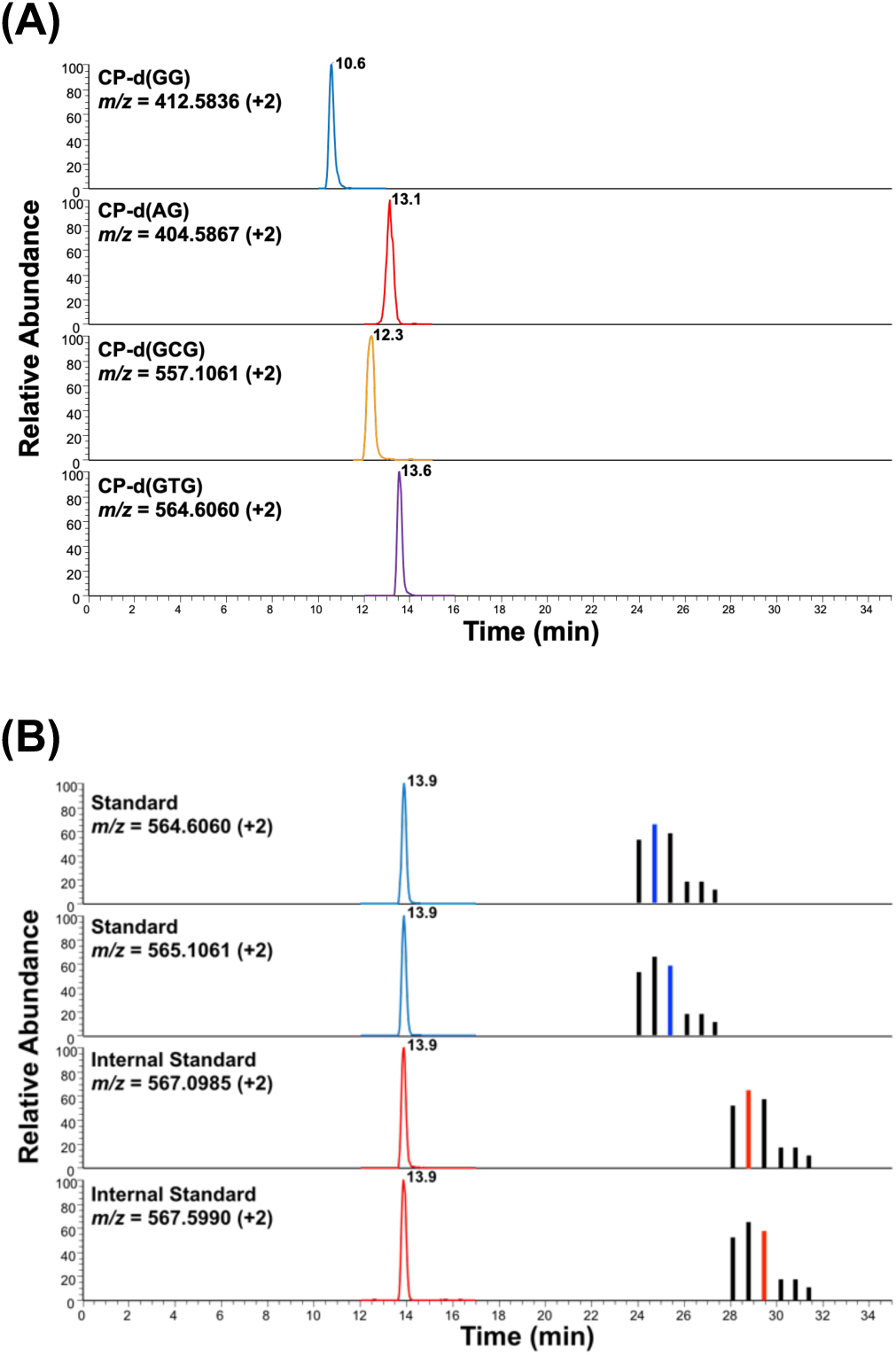
(**A**) Representative UPLC-HRAM-SIM chromatogram of the most abundant isotopes of doubly charged CP-d(GpX) analytes. (**B**) Representative UPLC-HRAM-SIM chromatogram of CP-d(GTG) from a 1 μM cisplatin-treated CTDNA sample. Panels 1 and 2 are isotopes of CP-d(GTG) analyte and panels 3 and 4 are the analogous isotopes of CP-d(GTG) IS.

To provide additional confidence that the desired CP-d(GpX) was detected, we chose to monitor two abundant isotopes. To avoid any potential, theoretical overlap of analyte and internal standard signals, the third isotope signal (^196^Pt) was used for quantitation and the more abundant second isotope signal (^195^Pt) was used for confirmation whenever possible (**Figure 3B**). Full scan-HRAM-MS assays failed to provide sufficient sensitivity due to high background. The use of selective ion monitoring drastically improved selectivity of each CP-d(GpX) and increased overall sensitivity to each CP-d(GpX). Using the conditions described above, we observed similar sensitivity with UPLC-PRM as compared to our final UPLC-SIM assays.

The developed UPLC-SIM assay was fully validated by processing and analyzing untreated CTDNA (25 µg, N = 3) spiked with an increasing amount of authentic standard (50 fmol – 1 pmol) and 1 pmol IS as described above. The observed ratio of analyte/IS was plotted against the expected analyte/IS ratio to confirm linearity (r^2^ = 0.99) and calculate accuracy and precision for all four CP-d(GpX). The limit of quantitation (LOQ) was set to be a signal-to-noise ratio >10, and the limit of detection (LOD) was set to be a signal-to-noise ratio >3. From the validation experiments, the following LOQs were calculated; CP-d(GG) LOQ = 20 fmol, CP-d(GCG) LOQ = 20 fmol, CP-d(GTG) LOQ = 10 fmol, CP-d(AG) LOQ = 50 fmol. The individual accuracy and precision calculations for each point of the validation curves are provided in **Table S4** and **Figure S4**.

### Digestion of platinated oligos

A key challenge in our approach is to develop a digestion protocol employing endo- and exonucleases capable of digesting platinated DNA up to the cross-link while avoiding over-digestion of the internal phosphodiester bonds. We first set out to identify exonucleases that fulfill this requirement on ssDNA oligonucleotides. We incubated a known amounts of 42mers containing a site-specific cisplatin cross-link with a series of endonucleases. After incubation, the digestion products were resolved by 20% urea gel and visualized by SYBR gold staining. Enzyme concentration and incubation times were optimized to maximize digestion efficiencies.

For both the 1,2- and 1,3-intrastrand cross-links, incubation of 100 pmol oligonucleotide with 40 mU of the 5’ – 3’ exonuclease phosphodiesterase II (PDE II) for 4 – 24 hours successfully digested up-to the platinum cross-link. Increasing enzyme concentration up to 80 mU did not yield noticeable over-digestion of the substrates. Similarly, incubation with 2 U of the 3’ – 5’ exonuclease I (Exo I) for 4 – 24 hours successfully digested 1,3-intrastrand and 1,2-AG-intrastrand cross-link oligos up to the platinum cross-link (**Figure S5A**). Surprisingly, the 1,2-GG-intrastrand cross-link oligo was only partially digested, with evidence of digestion being stalled before the platinum cross-link. The 3’ – 5’ exonuclease III (Exo III), which is more active on dsDNA substrates, only partially digested both 1,2- and 1,3-intrastrand cross-link oligos under every condition investigated (**Figure S5B**). The 3’ – 5’ exonuclease phosphodiesterase I (PDE I), 5’ – 3’ T7 exonuclease (T7 Exo), and bi-directional exonuclease V (Exo V) and exonuclease VII (Exo VII) failed to digest the ssDNA substrate under any of the conditions investigated. A summary of all digestion enzymes investigated is provided in **Table S5**.

To confirm the digestion efficiency results above, the digestions were repeated at a 1 nmol scale and the digestion products were purified by HPLC and analyzed by full-scan LC-MS. The desired 24mer (5’ – 3’ digestion) and 21mer (3’ – 5’) products eluted at 26.5 minutes (**Figure S6A**). The exact mass of the 24mer containing a 5’-platinum cross-link (1,2-GG *m/z* = 7407.1729 and 1,3-GTG *m/z* = 7401.1756) and the 21mer containing a 3’-platinum cross-link (1,2-GG *m/z* = 6205.9899, 1,3-GTG *m/z* = 6510.0359) was observed as the major product for the PDE II and Exo I digestions respectively (**Figure S6B & C**). As expected, the desired products were observed as a minor product for Exo III digestions (**Figure S6C**). Overall, our data show that PDE II, Exo I, and Exo III could appropriately digest platinated ssDNA oligonucleotides to yield CP-d(GpX).

### Digestion optimization of 1,2-intrastrand cross-links on calf thymus dsDNA substrates

We then set out to establish adequate digestion conditions of cisplatin adducts in dsDNA in the context of calf-thymus DNA (CTDNA) using a combination of endo- and exonucleases, focusing first on 1,2-intrastrand crosslinks. CTDNA is an equivalent substrate to isolated and deproteinized genomic DNA, and often used as an *in vitro* substrate to optimize DNA digestion conditions. Therefore, we treated CTDNA with an increasing concentration of activated cisplatin (50 nM – 10 µM) and performed digestions with multiple digestion methods. CP-d(GpX) were measured using the developed UPLC-SIM assays and isotope dilution of spiked isotopically labeled internal standards described above.

Previous attempts to quantitate cisplatin cross-links have utilized combinations of DNaseI, Nuclease P_1_ (NucP_1_), and alkaline phosphatase to digest platinated DNA to CP-d(GG)^21^. Analogously, Hah *et al* utilized a combination of DNaseI and Nuclease S_1_ (NucS_1_) to quantitate all possible oxaliplatin-induced DNA cross-links^23^. Finally, the commercially available Nucleoside Digestion Mix (New England Biolabs, Ipswich, MA) has been utilized to quantitate structurally diverse DNA adducts^26–27^. Given that all three methods utilize enzymes that are more efficient on dsDNA, we decided to include these conditions in our optimization experiments. As they yield digestion products that cannot be detected by urea gel analysis, these enzymes were not included in the previous analysis in Figure S4.

In agreement with the conditions published by Baskerville-Abraham *et al*^21^, sequential incubation with DNaseI, NucP_1_, and alkaline phosphatase (*Method 1*) yielded the highest levels of 1,2-GG-intrastrand cross-links. CP-d(GG) was detected at cisplatin treatments as low as 50 nM with 9.68 ± 0.81 CP-d(GG) per 10^6^ nucleotides (N = 3). Increasing cisplatin treatment of 100, 250, 500, and 1000 nM yielded a linear increase of 16.86 ± 0.54, 43.89 ± 3.1, 82.46 ± 8.4, and 146.52 ± 12.3 CP-d(GG) per 10^6^ nucleotides (N = 3, **Figure 4A**) respectively. Using *Method 1,* we observed significant suppression from 2’-deoxyadenosine that eluted at ∼17.3 minutes in HPLC and the same time on UPLC-SIM. To eliminate this from analysis, we modified *Method 1* by adding 0.005 U adenosine deaminase per 50 µg of CTDNA and incubating an additional 30 minutes at 37 °C. This modification resulted in a linear increase of 0.26 ± 0.12, 0.85 ± 0.03, 1.94 ± 0.13, 4.53 ± 0.18, and 10.04 ± 1.38 CP-d(AG) per 10^6^ nucleotides (N = 3) across 50 – 1000 nM cisplatin treatment respectively (**Figure 4A**).

**Figure 4:**
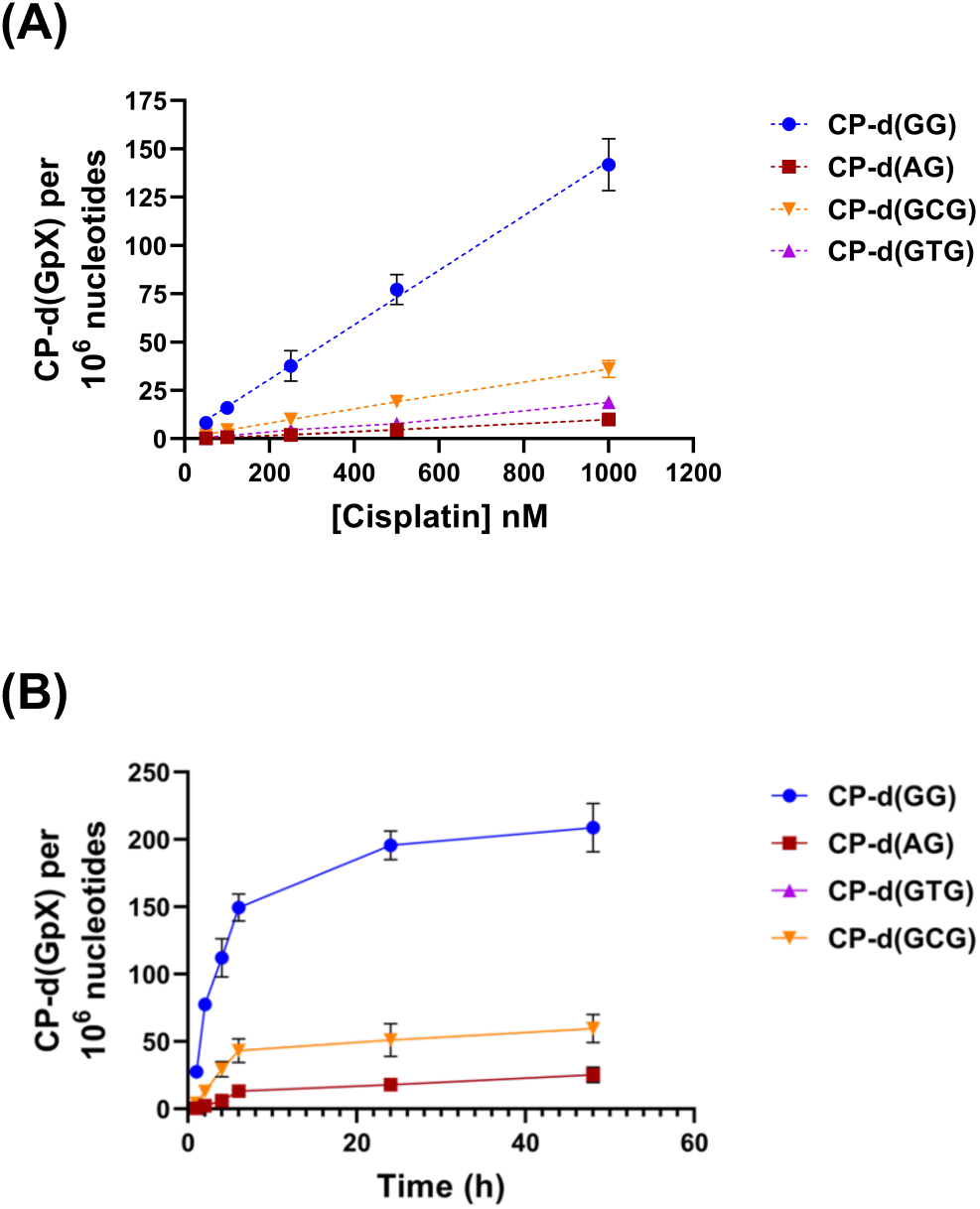
Quantitation of CP-d(GpX) of fully aquated cisplatin-treated CTDNA. Aliquots of platinated CTDNA (50 nM to 1 μM, N=3) were spiked with 1 or 2 pmol of the appropriate CP-d(GpX) IS, digested, and processed as described above depending on the adduct. Adduct levels are expressed as CP-d(GpX) per 10^6^ nucleotides. Linearity of the digestion protocols was confirmed by analyzing a dose-dependence set (50 nm – 1 μM) by the best available assay; Method #1 for CP-d(GG) and CP-d(AG), Method #3 for CP-d(GCG) and CP-d(GTG). (**B**) Intrastrand cross-link formation kinetics from CTDNA treated with cisplatin. CTDNA was incubated with 500 nM cisplatin for 1 – 48 hours. Aliquots of each time point (N = 3) were spiked with 1 or 2 pmol of the appropriate CP-d(GpX) IS, digestion, and enrichment as described above depending on the adduct. Adduct levels are expressed as CP-d(GpX) per 10^6^ nucleotides. Data points for CP-d(GTG) are behind data points of CP-d(AG).

For comparison, CP-d(GG) and CP-d(AG) levels were quantitated following the remaining digestion methods described above. *Method 4* (NucS_1_ 1 U per 5 µg CTDNA, ON incubation at 37 ^°^C) yielded the second-best results for 1,2-intrastrand cross-links with 58.18 ± 12.69, 114.41 ± 12.29, 278.81 ± 48.83, and 570.59 ± 88.83 CP-d(GG) per 10^6^ nucleotides (N = 3) and 1.48 ± 0.49, 2.43 ± 0.47, 6.54 ± 2.65, and 12.06 ± 2.42 CP-d(AG) per 10^6^ nucleotides (N = 3) for 1 – 10 µM CP treatment respectively (**Figure S7A**). *Method 3* (1 U NEB nucleoside digestion mix per 5 µg CTDNA, 2 hour incubation at 37 ^°^C) yielded the next best results for CP-d(GG) across the same CP range, but CP-d(AG) levels were below the LOQ and only detectable when the digestion time was increased to four hours (**Figure S7A and S7B**). Finally, *Method 2* was the least efficient for CP-d(GG), with adduct levels from 1 µM CP-treated CTDNA over 100-fold lower at 1.28 ± 0.27 adducts per 10^6^ nucleotides (N = 3, **Figure S7A**). Furthermore, quantitation across 1 – 10 µM CP treated CTDNA produced a non-linear response indicative of inconsistent digestion efficiency. Conversely, CP-d(AG) levels were below the LOQ for 50 – 500 nM CP but yielded a linear response across 1 µM – 10 µM CP (N = 2) treatment (**Figure S7B**). In conclusion, *Method 1* was clearly the most efficient at digesting the 1,2-intrastrand cross-links (**Table 2**).

**Table 2:** Comparison of CP-d(GpX) digestion efficiency for all investigated methods Comparison of CP-d(GpX) quantitation by all four assays from 1 μM cisplatin-treated CTDNA.

| Method | CP-d(GG) | CP-d(AG) | CP-d(GCG) | CP-d(GTG) |
| --- | --- | --- | --- | --- |
| 1<br>(DNase/NucP<sub>1</sub>) | 146.52 ± 12.3 | 9.57 ± 1.27 | <LOQ | <LOQ |
| 2<br>(DNase/Exo III/PDE II) | 1.28 ± 0.27 | 9.42 ± 2.49 | 1.18 ± 0.22 | 3.52 ± 0.93 |
| 3<br>(NEB nucleoside mix) | 27.07 | <LOQ | 36.18 | 18.83 |
| 4<br>(Benzonase/NucS<sub>1</sub>) | 54.18 ± 12.7 | 1.48 ± 0.49 | 6.48 ± 2.06 | 2.83 ± 0.53 |

### Digestion optimization of 1,3-intrastrand cross-links on dsDNA substrates

No methods that can accurately detect and quantitate 1,3-intrastrand cross-links have been reported to date, and we therefore attempted to optimize every developed digestion method described above on cisplatin treated CTDNA. The endonuclease NucP_1_ completely digested both CP-d(GCG) and CP-d(GTG) analyte and internal standard within one hour of digestion, including hydrolysis of the internal phosphodiester bonds, and *Method 1* could not be optimized to prevent over-digestion (data not shown). *Method 2* digestion yielded a broad CP-d(GCG) peak that eluted 0.1 minutes earlier than the internal standard (**Figure S8A**) and three different CP-d(GTG) analyte peaks at 13.6 minutes (small peak), 14.0 minutes (broad peak), and 14.5 minutes (small peak) across 1 – 10 µM CP treatment (**Figure S8B**). The same retention time shifts and multiple peaks were observed when *Method 3* and *Method 4* were initially investigated as well. Furthermore, over-digestion of the CP-d(GCG) and CP-d(GTG) IS was very inconsistent with both *Method 3* and *Method 4*.

Analysis of the raw data revealed that the shifted CP-d(GCG) analyte peak and all three CP-d(GTG) analyte peaks had the desired isotope distribution pattern and correct *m/z* ratios within 5 ppm, demonstrating that they represent CP-d(GCG) CP-d(GTG), respectively, possibly as a distinct rotamer or isomer. Heating the samples at 80 °C for 10 minutes prior to UPLC-SIM analysis did not yield a single peak with the same retention time as the internal standards, suggesting that the unexpected peaks may not be rotamers. UPLC-PRM analysis of the CP-d(GCG) and CP-d(GTG) analytes from method 4 digestion yielded the same major fragments from cleavage of the glycosidic bonds and one or two amine groups to yield two guanines cross-linked by platinum ([Gua–Pt(NH_3_)_x_– Gua]^+^ where X = 0, 1, or 2). However, the CP-d(GCG) analyte PRM yielded a unique fragment of 379.05571 which was identified as guanine cross-linked to platinum ([Gua– Pt(NH_3_)_2_]^+^), a unique fragment of 655.03366 which was not identified, failed to produce the fragment of guanine monophosphate-platinum-guanine base cross-link [GMP-Pt(NH_3_)-Gua]^+^, and the doubly-charged [Gua–Pt(NH_3_)_x_–Gua]^+^ fragments were less intense compared to authentic standards and internal standards. Furthermore, the CP-d(GTG) PRM of peaks 2 and 3 yielded a unique fragment of 789.05365 which was not identified. Overall, the data confirmed that the 1,3-intrastrand cross-link analytes identified contained a GCG or GTG trimer cross-linked between the terminal guanines, and they most likely constitute structural isomers of the expected N7-N7 cross-linked CP-d(GXG).

To elucidate the etiology of these unexpected peaks, we investigated whether they were present after digesting either single-stranded or short double-stranded cisplatin cross-link substrates. Digestion of 100 pmol 42mer single-stranded oligonucleotide containing a site-specific 1,3-GCG or 1,3-GTG cisplatin crosslink by Method 3 yielded only one CP-d(GCG) or CP-d(GTG) analyte peak with the same retention time as the internal standard. To investigate a simple double-stranded substrate with one possible 1,3-intrastrand cross-link position, the 42mer 1,3-GTG oligo substrate was annealed with a complementary strand, followed by incubation with cisplatin (1:1, 1:3, or 1:10 dsDNA:CP ratio). Digestion of this substrate once again yielded a single CP-d(GTG) analyte peak with the same retention time of the internal standard (**Figure S8C**). Based on these experiments, we conclude that the unexpected peaks are not artifacts of enzymatic digestion and must originate from secondary structures of complex double-stranded DNA and could involve isomerization following formation of the original crosslink.

Since each observed analyte peak were confirmed to be an isomer of the desired CP-d(GCG) and CP-d(GTG), we took the combined peaks as a measure for the 1,3 intrastrand adducts for the evaluation of the remaining digestion methods. Under the initial digestion conditions, both the NEB nucleoside digestion mix of method 3 and NucS_1_ of Method 4 completely over-digested the internal standard, corresponding analyte peaks, and peak 3 of CP-d(GTG) analyte. Furthermore, the remaining CP-d(GCG) analyte and peak 2 of CP-d(GTG) analyte were also partially digested. To reduce undesired over-digestion, the digestion conditions and incubation times were further optimized.

When NucS_1_ concentration was decreased to 1 U per 10 µg CTDNA and incubation time decreased to 1 hour at 37 °C, over-digestion was decreased to ∼10% for both CP-d(GCG) and CP-d(GTG). Using these new NucS_1_ digestion conditions, a linear increase of 7.45 ± 1.49, 14.16 ± 1.35, 34.27 ± 4.66, and 67.87 ± 10.63 CP-d(GCG) per 10^6^ nucleotides and 17.70 ± 6.75, 33.72 ± 11.83, 84.22 ± 28.15, and 170.54 ± 6.38 CP-d(GTG) per 10^6^ nucleotides across 1, 2, 5, and 10 µM CP respectively (N = 3) was observed.

The NEB nucleoside digestion mix was optimized by incubating 1 pmol of CP-d(GXG) standard and IS with a decreasing amount of enzyme (2.5 U, 0.5 U, 0.25 U, and 0.05 U) over different periods of time (1, 5, 10, 30, and 60 minutes). CP-d(GCG) was immediately digested with the original conditions, with only 57, 39, 28, 8.4, and 1.2% remaining after 1, 5, 10, 30, and 60 minutes respectively (**Figure S9A**). CP-d(GTG) was more resistant to over-digestion, with 90, 72, 62, 50, and 40% remaining across the same time points (**Figure S9B**). Based on these initial results in solution, we decided to investigate over-digestion of standard and IS spiked into 25 µg of CTDNA with 2.5, 1.0, 0.75, and 0.5 U of NEB nucleoside digestion mix over 2, 10, and 30 minutes (**Figure S9C**). Overall, we decided to use 1.0 U NEB nucleoside digestion mix per 25 µg DNA for 30 minutes (∼40% over-digestion) due to efficient CTDNA digestion and the inconsistency observed with the original conditions. There was no clear trend for CP-d(GTG) (∼20 to 30% over-digestion, **Figure S9D**), allowing us to use the same conditions as CP-d(GCG).

Utilizing the new *Method 3* conditions, the overall IS signal was about four-fold greater. A linear increase of 1.97 ± 0.95, 4.39 ± 0.98, 10.0 ± 1.5, 19.24 ± 2.8, and 36.18 ± 4.59 CP-d(GCG) per 10^6^ nucleotides and 0.95 ± 0.41, 1.56 ± 0.50, 4.61 ± 1.06, 7.76 ± 0.67, and 18.83 ± 2.87 CP-d(GTG) per 10^6^ nucleotides across 50 – 1000 nM CP respectively (N = 3, **Figure 4A**). *Method 2* was nearly six-fold less efficient for both CP-d(GCG) and CP-d(GTG) but did yield a linear increase across 1 – 10 µM CP treatment (N = 2). Based on the results above, we decided to only utilize the optimized *Method 3* with for CP-d(GXG).

### CP-d(GpX) formation kinetics in CTDNA

To better understand intrastrand cross-link formation from cisplatin and further validate our methodology, we quantified intrastrand cross-link formation over time. CTDNA was incubated with 500 nM cisplatin and samples were quenched with 10 mM thiourea to prevent further adduct formation during the work up and digestion. Samples were processed at various time points from 1 – 48 hours (N = 3). The 1,2-GG-intrastrand cross-links formed at the fastest rate with 27.63 ± 2.3, 77.67 ± 2.6, 112.09 ± 11.6, and 149.49 ± 8.17 CP-d(GG) per 10^6^ nucleotides after 1, 2, 4, and 8 hours of cisplatin incubation respectively (**Figure 4B**). The levels then plateaued at 195.81 ± 8.8 and 208.86 ± 14.6 CP-d(GG) per 10^6^ nucleotides across 24 and 48 hours respectively. The 1,3-GCG-intrastrand cross-links formed at the next fastest rate with 4.2 ± 0.32, 12.99 ± 1.8, 29.50 ± 4.7, 43.16 ± 7.2, 51.19 ± 9.9, and 59.71 ± 8.6 CP-d(GCG) per 10^6^ nucleotides in the same time frame respectively. The 1,3-GTG-intrastrand cross-links formed faster initially with 1.81 ± 0.2, 5.39 ± 0.4, and 12.55 ± 2.4 CP-d(GTG) per 10^6^ nucleotides across 1, 2, and 4 hours respectively, but more quickly plateaued with 17.22 ± 1.2, 21.74 ± 3.7, and 27 ± 4.8 CP-d(GTG) per 10^6^ nucleotides across 8, 24, and 48 hours respectively. Finally, 1,2-AG-intrastrand cross-link formation showed a similar trend to 1,2-GG and 1,3-GTG with 0.62 ± 0.33, 2.49 ± 0.34, 6.15 ± 1.5, and 13.15 ± 3.4 CP-d(AG) per 10^6^ nucleotides across 1, 2, 4, and 8 hours respectively, followed by plateauing with 18.01 ±1.6 and 25.41 ±4.7 CP-d(AG) per 10^6^ nucleotides across 24 and 48 hours respectively.

Using the data from 48 hour cisplatin exposure, the new adduct distribution is 65.4%, 8.0%, 18.7%, and 7.9% for 1,2-GG, 1,2-AG, 1,3-GCG, and 1,3-GTG-intrastrand cross-links respectively. Taking into account over-digestion of 1,3-intrastrand cross-links, the distribution is 71.2%, 8.7%, 13.2%, and 6.9% for 1,2-GG, 1,2-AG, 1,3-GCG, and 1,3-GTG-intrastrand cross-links respectively. Based on previously reported cross-link distributions^5, 8^, the methods for 1,2-GG and 1,3-intrastrand cross-links were completely validated and the method for 1,2-AG-intrastrand cross-links requires further optimization.

### CP-d(GpX) quantitation in CP-treated cell lines

To demonstrate the applicability of our developed assays to DNA repair and biomonitoring experiments in cells, we quantitated CP-d(GpX) repair over time in CP-treated isogenic U2OS WT and XPA-deficient (NER-deficient) cells. Both cell lines were initially treated with 10 µM CP for 2 hours, followed by removal of the drug was removed and allowing to recover for 0, 1, 2, 4, 8, and 24 hours. Cells were then lysed, genomic DNA extracted and purified and the extracted DNA from a single 10 cm dish (N = 3) was digested by *Method 1* and *Method 3* quantitate CP-d(GG) and CP-d(GXG) analytes, respectively. These preliminary results revealed that both CP-d(GCG) and CP-d(GTG) levels were near the LOQ, leading to inconsistent quantification. We therefore decided to repeat the treatment with 25 µM CP following the same exact procedure above.

The most abundant 1,2-GG-intrastrand cross-link was repaired very slowly in NER-proficient cells, with 50% of initial adduct remaining after 8 hrs incubation, but the levels did not decrease over the same time period in the NER-deficient cell lines (**Figure 5A**). The 1,3-intrastrand cross-links were measured from 25 µM CP-treated U2OS cells. CP-d(GCG) levels decreased slowly in NER-proficient cells similar to CP-d(GG), with about 60% of initial adduct levels remaining after 8 hrs. However, CP-d(GCG) levels were significantly higher and remained constant over an 8 hr time period in NER-deficient cells (**Figure 5B**). By comparison, CP-d(GTG) had a similar trend in NER-proficient cells as for the CP-d(GG) adducts, with about 45% of adducts remaining after 8 hrs, while the adduct levels remained constant over the same time period in NER-deficient U2OS cells. Taken together, our assays were able to quantitate the repair of 1,3-intrastrand cross-links in NER-proficient cell lines and the accumulation of 1,2- and 1,3-intrastrand cross-links in NER-proficient cell lines, demonstrating their applicability for future kinetic and mechanistic studies of DNA repair of cisplatin DNA intrastrand crosslinks.

**Figure 5:**
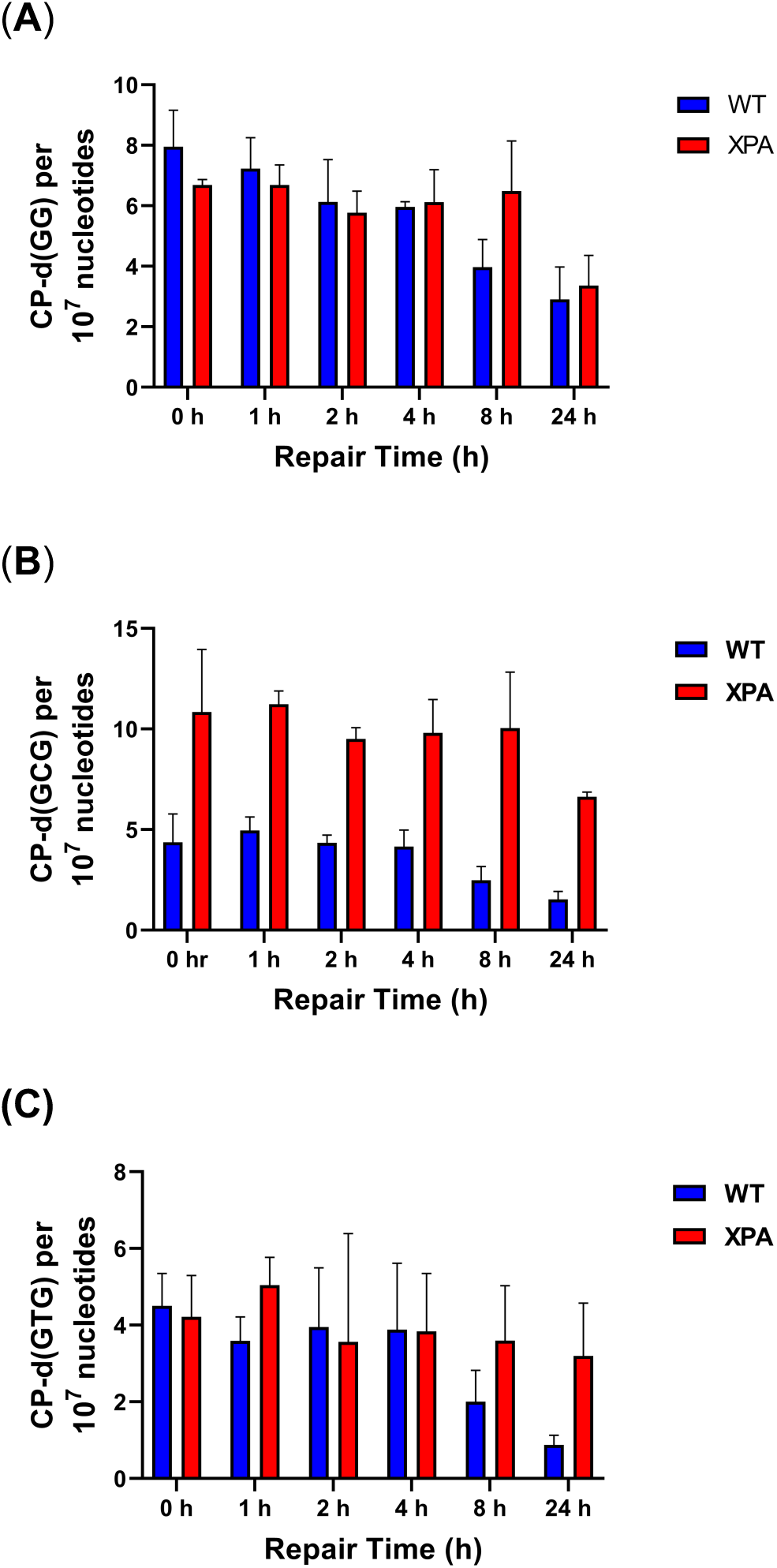
Quantification of CP-d(GpX) in cisplatin-treated NER-deficient cells. Aliquots of platinated genomic DNA (2 h incubation with 10 or 25 μM cisplatin, 0, 1, 2, 4, 8, and 24 h repair time post treatment, N = 3) were spiked with 1 pmol of appropriate CP-d(GpX) IS and processed as described above depending on the adduct. Total CP-d(GpX) adducts were quantitated by UPLC-SIM for (**A**) CP-d(GG) and CP-d(AG) obtained by method 1 and (**B**) CP-d(GCG) and CP-d(GTG) obtained by method 3.

## DISCUSSION

Cisplatin is one of the most widely used chemotherapeutic agents to treat solid tumors, with a cure rate of over 90% for testicular cancers. Unfortunately, cisplatin treatment often yields toxic side effects including nephrotoxicity and ototoxicity, and many patients develop resistances to the treatment^1-2, 4^. Although the exact mechanism(s) of these side effects are multifaceted, the formation, recognition, and repair of DNA-DNA cross-links is obliviously a crucial factor. While methods have been reported for the analysis of the most abundant cisplatin adduct, the 1,2-GG-intrastrand cross-link, have been reported, current methods do not allow for the quantification of the remaining adducts formed. This lack of information makes it impossible to fully elucidate how much each of the DNA adducts contributes to the toxicity of cisplatin and how the formation, recognition, and repair of DNA damage contributes to drug resistance.

To address this need, we have developed a comprehensive UPLC-HRAM-SIM assay to measure 1,2- and 1,3-intrastrand cross-links. Similar to previously developed LC-MS/MS^21^ and ICP-MS^20^ assays for the most abundant 1,2-GG-intrastrand cross-link, we were able to develop a CP-d(GG) assay with a LOQ of 20 fmol, or 10 1,2 CP-d(GG) per 10^8^ nucleotides. In addition, we have developed and fully validated the first isotope dilution mass spectrometry-based assays to quantitate 1,2-AG-intrastrand, 1,3-GCG-intrastrand, and 1,3-GTG-intrastrand cross-links with LOQs of 50 fmol, 10 fmol, and 10 fmol, respectively.

While optimizing the digestion methodologies, we observed a shift in the CP-d(GCG) analyte and multiple CP-d(GTG) peaks from CP-treated CTDNA compared to the standard generated from treatment of nucleotide trimers with cisplatin. We do not currently know whether these are chemical isomers (different connectivity of the cisplatin and DNA base) or structural isomers (for example rotamers). Coordination between platinum and the phosphodiester bonds of DNA has been proposed^28^, but UPLC-PRM analysis confirmed that these peaks were the desired trimer with platinum cross-link between the terminal guanines. Although heating at 80 °C for 10 minutes did not reverse any peaks, it is possible that the observed shifted and extra peaks are extremely stable rotamers formed after release from DNA. The observation of a single analyte peak for CP-d(GG) and CP-d(AG) is in agreement with an x-ray crystal structure of 5’-dCGG cross-link which showed no steric rotation^29^. Furthermore, it is possible that one cross-linking position between GNG sites can occur at another nucleophilic sites such N1 or O^6^ of guanine, yielding chemical isomers with different elution times compared to the internal standards. However, considering that all of them have the exact mass and isotope distribution of the 1,3 intrastrand crosslinks, we conclude that they are a reliable measure of cisplatin adduct levels.

Quantitation of all CP-d(GpX), including the shifted CP-d(GCG) analyte peak and the multiple CP-d(GTG) analyte peaks, yielded a dose-dependent linear response (**Figure 4A**), further validating the optimized methodologies can be used to quantify intrastrand cross-links. Using the results from 1 µM CP (**Table 2**), we observed an overall adduct distribution of 68.6% 1,2-GG, 4.8% 1,2-AG, 17.5% 1,3-GCG, and 9.1% 1,3-GTG. Accounting for over-digestion of CP-d(GCG) and CP-d(GTG), the adduct distribution is 74.5% 1,2-GG, 5.3% 1,2-AG, 12.3% 1,3-GCG, and 7.9% 1,3-GTG or 79.8% 1,2-intrastrand and 20.2% 1,3-intrastrand cross-link. In comparison to previous studies, our quantified 1,2-AG adduct levels were nearly 5-fold lower and those of the 1,3-intrastrand cross-link 2-fold higher than expected^6-8^.

These results could be explained by our use of fully activated cisplatin Pt(NH_3_)_2_(H_2_O)_2_, which alkylates both the 5’ and 3’ positions of -GG-sequence^30–31^ and the 5’ position of -GA-sequence much faster than cisplatin^32^. Therefore, the rapid accumulation of monoadducts and CP-induced cross-links will create segments of ssDNA more accommodating to 1,3-intrastrand and interstrand cross-linking. In agreement with this hypothesis, Eastman observed a 2-fold decrease in 1,2-AG-intrastrand cross-links, 3-fold increase of interstrand cross-links, and near 10-fold increase in “undigested” DNA when CTDNA was denatured before platination^8^. Furthermore, the levels of 1,3-intrastrand and interstrand cross-links may have been underestimated because of incomplete digestion of 1,3-intrastrand cross-links by either nuclease P_1_ or nuclease S_1_, yielding a dG-cis-dG digestion product with 0, 1, or 2 phosphate groups^33^.

To further evaluate cross-link distribution from cisplatin, we quantified CP-d(GpX) formation kinetics over a 1 – 48 hour period. This would allow us to better compare our adduct distribution to previous results and ensure that 1,2-AG-intrastrand cross-links did not degrade or convert to another cross-link over time. As demonstrated in **Figure 4B**, 1,2-GG-intrastrand cross-links formed rapidly in the first 4 hours before plateauing at 24 and 48 hours. In comparison, 1,2-AG-intrastrand cross-links formed at a slower pace but also plateaued across 24 and 48 hours. This cross-link kinetics profile is in agreement with previous *in vitro* experiments showing that 1,2-GG-intrastrand cross-linking was much faster than other 1,2-intrastrand cross-links^31–32^. Our analysis of 1,3-GCG and 1,3-GTG-intrastrand cross-linking is the first kinetic formation data for 1,3-intrastrand cross-links and is in agreement with 1,3-intrastrand cross-links forming slower than 1,2-GG-intrastrand cross-links.

There is evidence that the 1,2-AG-intrastrand cross-links are more difficult to digest compared to 1,2-GG-intrastrand cross-links^6, 34^. It is possible that our digestion methodology could be further specifically optimized for 1,2-AG-intrastrand cross-links. Additionally, the cross-link between N7 of adenine and N7 of guanine may be less stable than between two guanines^35^, and isolated CP-d(AG) may therefore degrade during our sample enrichment. Isomerization of the ligands on platinum from cis to trans is not considered a possibility since transplatin does not readily form 1,2-intrastrand cross-links^36^. Overall, obtaining comprehensive kinetics of CP-induced DNA-DNA cross-linking and how adduct profiles change in different environments are future goals which our assays are uniquely qualified to address.

Detailed kinetic and molecular modeling studies to determine the sequence preference of CP to 5’-GG-3’ and 5’-AG-3’ compared to 5’-GA-3’ have yielded conflicting results, further complicating our understanding of the potential relevance of the different intrastrand cross-links. [H^1^,N^15^] HSQC 2D NMR time-course studies revealed that the rate of monoalkylation on the 3’-adenosine of 5’-GA-3’ by CP is 80 – 200 fold slower than alkylation of the 3’-position of either 5’-AG-3’ and 5’-GG-3’ respectively, and ring closure of 5’-AG-3’ from a 3’-alkylated-G is nearly 5 and 14-times faster than ring closure of 5’-GA-3’ from a 3’-alkylated-A and 5’-alkylated-G respectively ^32^. If cisplatin is replaced with fully activated cisplatin, the rate of monoalkylation is similar but ring closure of 5’-AG-3’ is nearly 23-fold faster. Molecular modeling studies revealed activation energies of 23 and 32 kcal/mol for 5’-AG-3’ and 5’-GA-3’ respectively, suggesting 5’-AG-3’ cross-linking would be 6-fold faster^37^. Taken together, a clear preference for 5’-AG-3’ over 5’-GA-3’ cross-link formation is favored. However, Monjardet-Bas *et al*^38^ observed only a two-fold difference in reactivity of CP between 5’-GA-3’ and 5’-AG-3’ sequences in the same double-stranded oligo, and Gupta *et al*^39^ confirmed cross-linking at 5’-GA-3’ even in the presence of more reactive sequences. We observed clear evidence of CP-d(GA) after total digestion of CTDNA treated with aquated CP, providing the first direct evidence of 1,2-GA-intrastrand cross-linking in complex double-stranded DNA and confirming cross-linking in the presence of multiple competing positions. However, the levels were significantly lower than any other intrastrand cross-link investigated, agreeing with a strong preference for 5’-AG-3’ over 5’-GA-3’ sequences.

We observed slow repair of both 1,2- and 1,3-intrastrand cross-links in CP-treated U2OS wild type cells, but no evidence of repair for intrastrand crosslinks in CP-treated U2OS XPA^-/-^ cells (**Figure 5A – 5C**). This is in complete agreement with the primary repair mechanism of CP-induced intrastrand cross-links being nucleotide excision repair^40–44^. The slow repair of 1,2-intrastrand cross-links has been observed with analogous ICP-MS^45–46^ and immunological assays^47^, although faster, biphasic repair kinetics was observed with early experiments^48–49^. Many transcription factors such as HMG proteins^50–52^, Ixr1^53–54^, mtTFA^51^, LEF−1^51^, and hUBF^55–56^, and mismatch repair proteins^57–59^ bind to 1,2-intrastrand cross-links. This could potentially inhibit the recruitment of NER factors and the efficient repair of 1,2-intrastrand cross-links. The slow repair of 1,3-intrastrand cross-links could be a product of the slow formation kinetics (**Figure 4B**), and the cross-links being repaired quickly after their formation.

In conclusion, we have developed sensitive and robust UPLC-SIM assays for the quantitation of cisplatin-induced 1,2-AG, 1,2-GG, 1,3-GCG, and 1,3-GTG intrastrand cross-links. Each individual assay was fully validated using *in vitro* samples and further utilized to measure CP-d(GpX) formation and repair in NER-deficient or isogenic control cell lines. We were able to observe the active repair of 1,3-intrastrand cross-links over time in NER-proficient cells and the persistence of 1,3-intrastrand cross-links in NER-deficient cells, demonstrating the utility of our assay for the study of repair kinetics. We are currently developing an analogous assay for oxaliplatin-induced cross-links to allow the quantitation of DNA-DNA cross-links from every FDA approved platinum-based drug. In the future, we hope to directly compare the formation and repair of platinum drug-induced DNA-DNA cross-linking *in vitro* and *in vivo*.

## FUNDING

This work was supported by the Korean Institute for Basic Science (IBS-R022-A1)

## ACKNOWLEDGMENTS

We thank Martijn Luijsterburg (University of Leiden) for the gift of WT and XPA-deficient U2OS cells.

## ABBREVIATIONS

(ACN): Acetonitrile
(AAS): Atomic absorption spectroscopy
(AGC): Automatic gain control
(CTDNA): Calf thymus DNA
(CP): Cisplatin
(CV): Column volumes
(DMEM): Dulbeccos modified eagles medium
(dsDNA): Double stranded DNA
(EtOH): Ethanol
(ESI): Electrospray ionization
(Exo I): Exonuclease I
(Exo III): Exonuclease III
(Exo V): Exonuclease V
(Exo VII): Exonuclease VIII
(FPLC): Fast protein liquid chromatography
(HPLC): High performance liquid chromatography
(HPLC-MS/MS): High performance liquid chromatography – tandem mass spectrometry
(HRAM): High resolution accurate mass
(ICP-MS): Inductively coupled plasma mass spectrometry
(IS): Internal standard
(ICL): Interstrand cross-link
(kDa): kilodalton
(LCMS): Liquid chromatography – mass spectrometry
(MS): Mass spectrometry
(µg): microgram
(mM): microliter
(µL): milligram
(mg): milliliter
(mL): millimolar
(mMol): millimole
(NEB): New England Biolabs
(NucP_1_): Nuclease P_1_
(NucS_1_): Nuclease S_1_
(NER): Nucleotide excision repair
(Pt): Platinum
(Quick CIP): Quick calf intestinal alkaline phosphatase
(rcf): Relative centrifugal force
(NaCl): Sodium chloride
(NaClO_4_): Sodium perchlorate
(TBE): Tris-borate-EDTA
(UPLC-PRM): Ultra performance liquid chromatography – parallel reaction monitoring
(UPLC-SIM): Ultra performance liquid chromatography – selective ion monitoring
(UP): Ultrapure
(UV): Ultraviolet

## Notes

### Competing Interest Statement

The authors have declared no competing interest.

