## Supplementary Materials for "Development of Comprehensive Ultraperformance Liquid Chromatography-High Resolution Mass Spectrometry Assays to Quantitate Cisplatin-Induced DNA-DNA Cross-Links"

### TABLE OF CONTENTS

Figure S1. Isotope distribution of CP-d(GpX) 2
Table S1. Table of observed CP-d(GpX) authentic standard fragments from UPLC-PRM experiments 3
Figure S2. CP-d(GpX) authentic standard fragmentation by UPLC-PRM 4
Table S2. Table of observed CP-d(GpX) internal standard fragments from UPLC-PRM experiments 5
Figure S3. CP-d(GpX) internal standard fragmentation by UPLC-PRM 6
Table S3. Effect of pH on CP-d(GpX) signal intensity 7
Table S4. Intraday validation of developed CP-d(GpX) assays 8
Figure S4. Intraday validation of developed CP-d(GpX) assays figures 9
Figure S5. Optimizing digestion efficiency to obtain CP-d(GpX) 10
Table S5. Summary of single-enzyme digestions of a 42mer cross-link substrate 11
Figure S6. Detailed analysis of 42mer cross-link substrate digestion by a single enzyme 12
Figure S7. Evaluating alternative digestion methods to quantify 1,2-intrastrand cross-links 13
Figure S8. Digestion of 1,3-intrastrand cross-links to CP-d(GXG) 14
Figure S9. Optimization of CP-d(GXG) digestion by Method #3 15

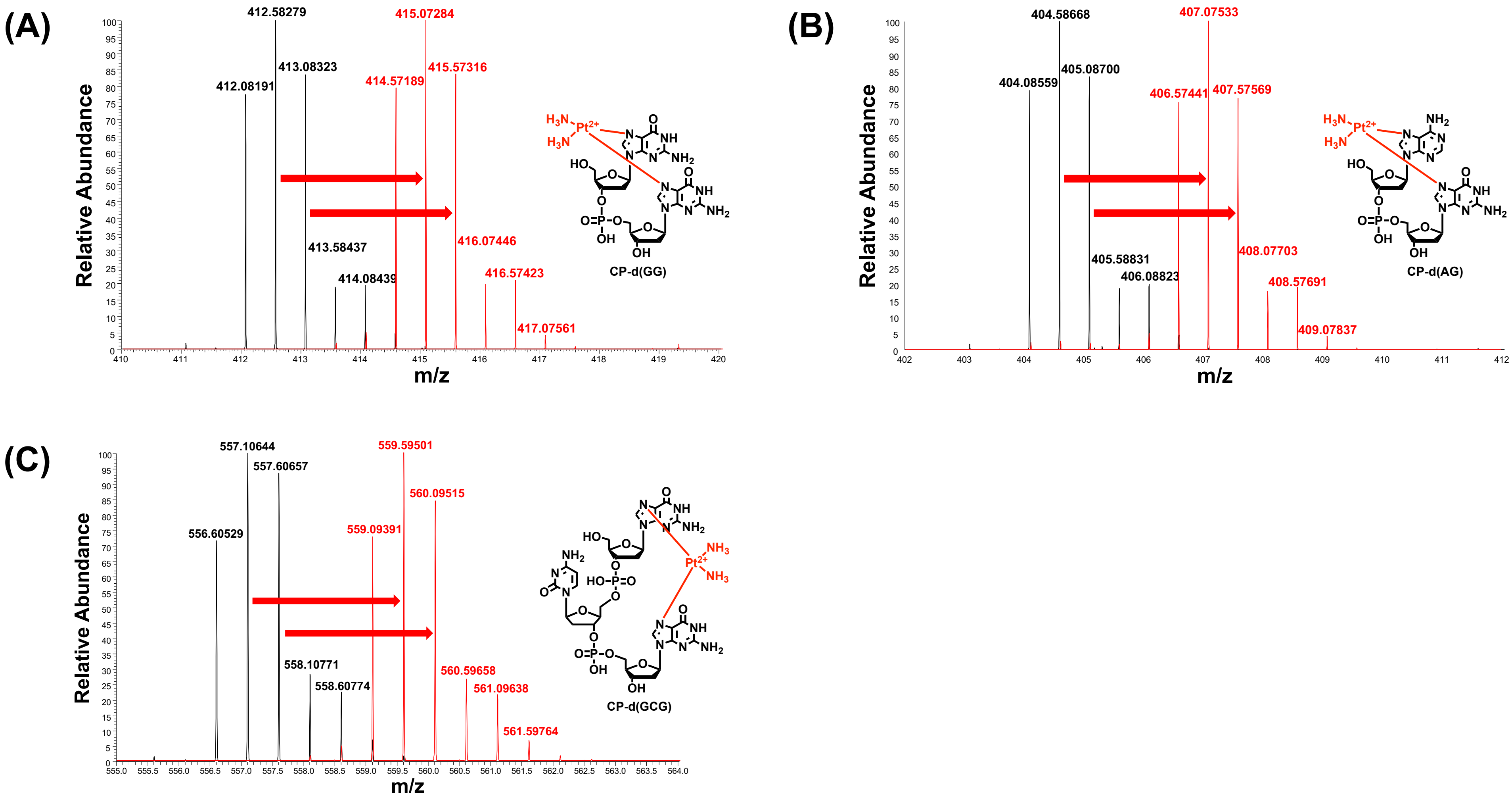


**Figure S1**: Isotope distribution of authentic standards and internal standards of **(A)** CP-d(GG), **(B)** CP-d(AG), and **(C)** CP-d(GCG).

| Substrate | Parent mass (m/z) | Observed fragment 1 (m/z) | Observed fragment 2 (m/z) | Observed fragment 3 (m/z) |
|---|---|---|---|---|
| CP-d(GA) | 404.0842 | | 479.0577 | 240.0324 |
| | 404.5851 | | 480.0598 | 240.5335 |
| | 405.0854 | | 481.0600 | 241.0336 |
| CP-d(AG) | 404.0900 | 496.0847 | 479.0582 | 248.5460 |
| | 404.5900 | 497.0866 | 480.0603 | 249.0470 |
| | 405.0900 | 498.0874 | 481.0608 | 249.5472 |
| CP-d(GG) | 412.0818 | 512.0796 | 495.0527 | 248.0299 |
| | 412.5827 | 513.0815 | 496.0548 | 248.5310 |
| | 413.0830 | 514.0818 | 497.0551 | 249.0312 |
| CP-d(GCG) | 556.6048 | 512.0794 | 495.0526 | 529.1058 |
| | 557.1057 | 513.0813 | 496.0543 | 530.1079 |
| | 557.6059 | 514.0815 | 497.0552 | 531.1080 |
| CP-d(GTG) | 564.1047 | 512.0795 | 495.0526 | 529.1061 |
| | 564.6060 | 513.0815 | 496.0546 | 530.1081 |
| | 565.1100 | 514.0817 | 497.0550 | 531.1084 |

**Table S1**: Fragmentation data is from the most abundant isotope. For all CP-d(GpX), the key fragments of dG-$Pt(NH_3)_2^+$, Gua-$Pt(NH_3)_2$-$Gua^+$, and Gua-$Pt(NH_3)$-$Gua^+$ were observed for every isotopes investigated.

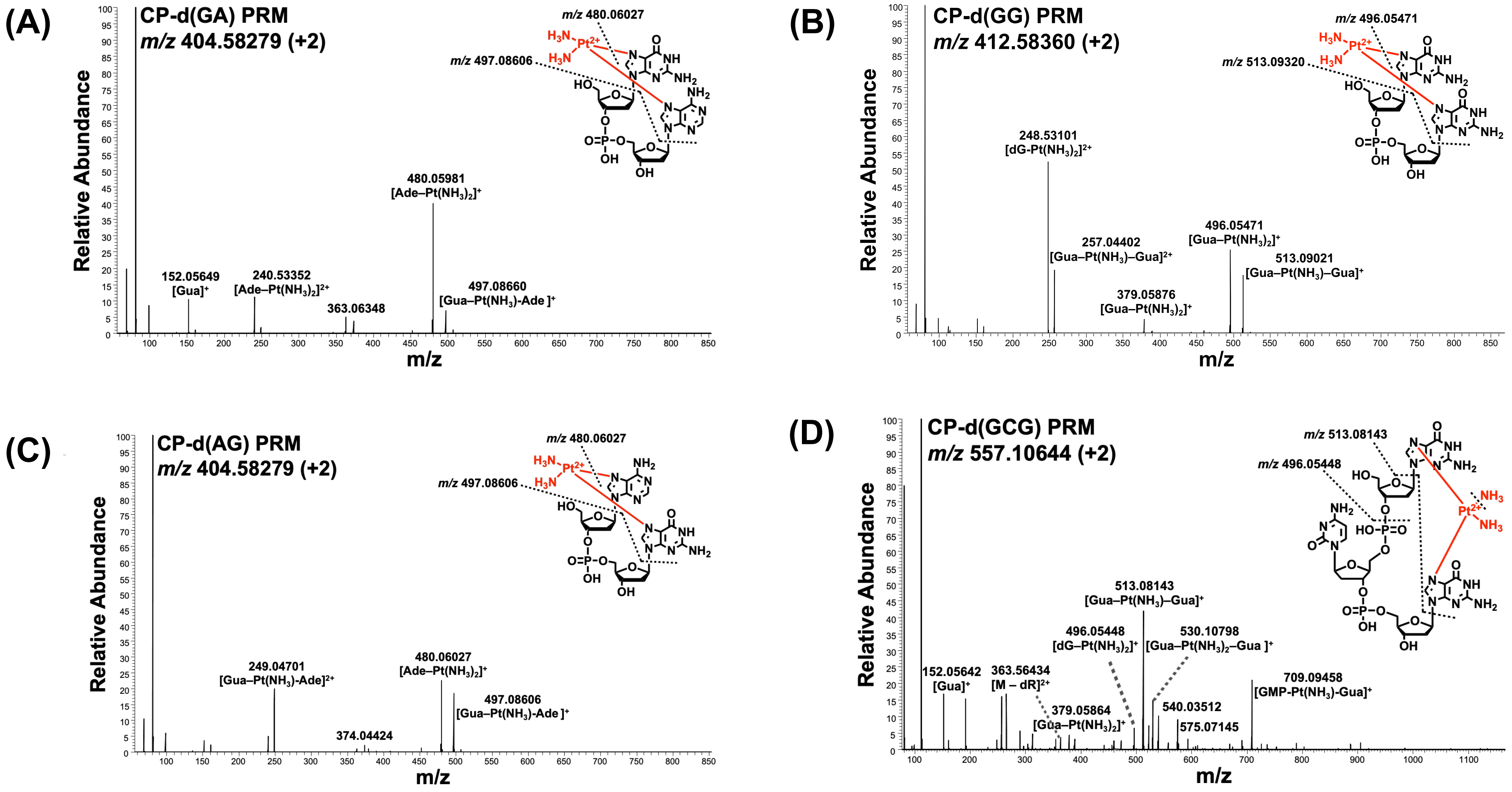


**Figure S2**: Representative traces of UPLC-PRM results for **(A)** CP-d(GA), **(B)** CP-d(GG), **(C)** CP-d(AG), and **(D)** CP-d(GCG) standards. Fragmentation data is from the most abundant isotope.

| Substrate | Parent mass (m/z) | Observed fragment 1 (m/z) | Observed fragment 2 (m/z) | Observed fragment 3 (m/z) |
|---|---|---|---|---|
| CP-d(AG) | 406.5745 | 484.0402 | 501.0666 | 251.0370 |
| | 407.0754 | 485.0421 | 502.0686 | 251.5379 |
| | 407.5757 | 486.0423 | 503.0689 | 252.0381 |
| CP-d(GG) | 414.5719 | 517.0615 | 500.0348 | 250.5211 |
| | 415.0729 | 518.0632 | 501.0371 | 251.0221 |
| | 415.5732 | 519.0641 | 502.0375 | 251.5223 |
| CP-d(GCG) | 559.0939 | 517.0614 | 500.0345 | 534.0881 |
| | 559.5949 | 518.0636 | 501.0368 | 535.0902 |
| | 560.0951 | 519.0637 | 502.0371 | 536.0905 |
| CP-d(GTG) | 566.5938 | 517.0614 | 500.0347 | 534.0879 |
| | 567.0946 | 518.0636 | 501.0367 | 535.0901 |
| | 567.5949 | 519.0638 | 502.0372 | 536.0902 |

**Table S1**: For all CP-d(GpX) internal standards, the key fragments of dG-Pt$(NH_3)_2^+$, Gua-Pt$(NH_3)_2$-Gua$^+$, and Gua-Pt$(NH_3)$-Gua$^+$ were observed for all isotopes investigated.

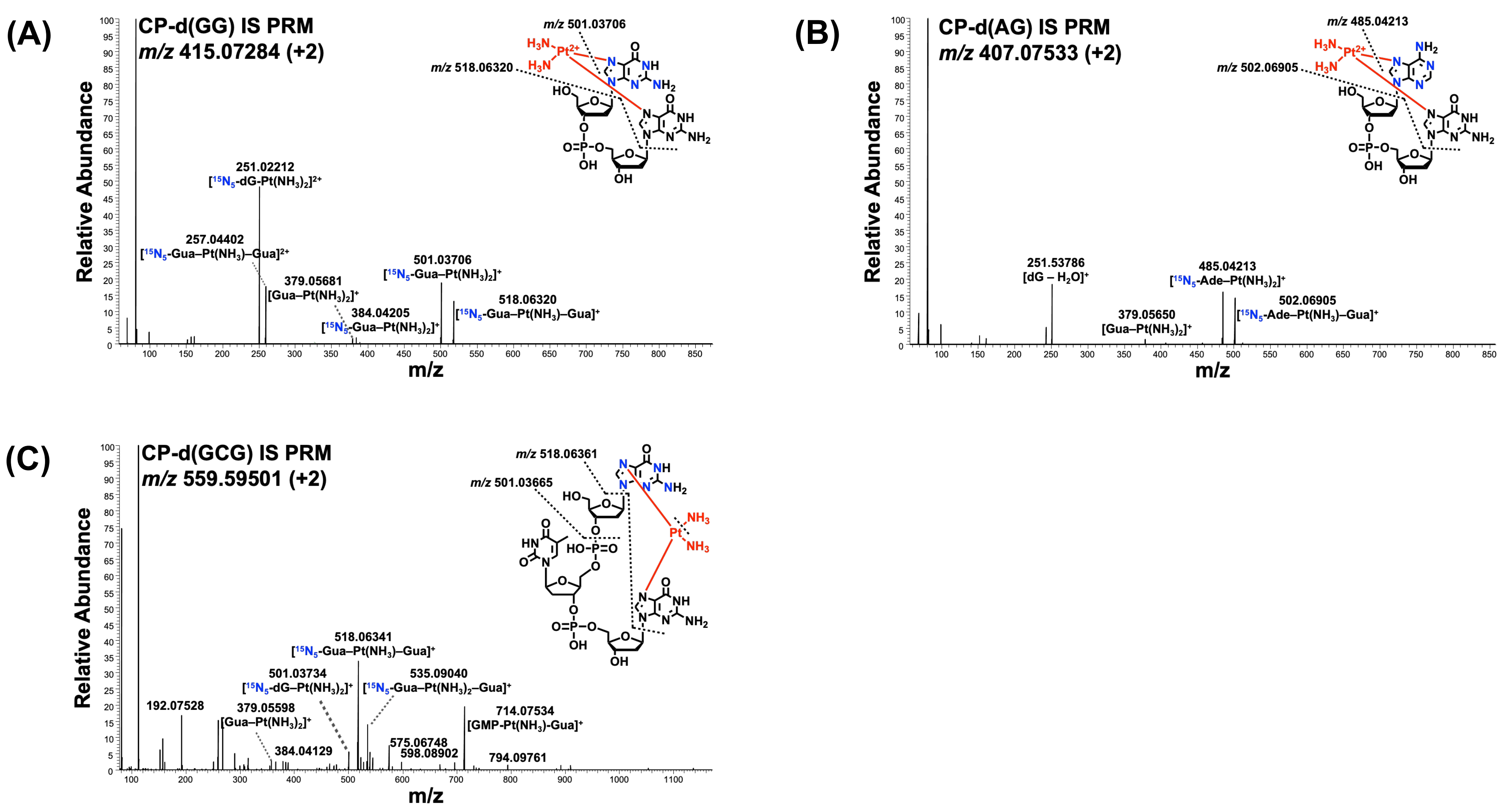


**Figure S3**: Representative traces of UPLC-PRM results for **(A)** CP-d(GG), **(B)** CP-d(AG), and **(C)** CP-d(GCG) internal standards. Fragmentation data is from the most abundant isotope.

| Analyte | pH 7.0 (Analyte area) | pH 5.5 (Analyte area) | pH 4.0 (Analyte area) | Comment |
| --- | --- | --- | --- | --- |
| CP-d(GG) | 1472816 | 1185339 | 792954 | Sensitivity decreased 46.2% |
| CP-d(AG) | 324120 | 311337 | 190041 | Sensitivity decreased 41.4% |
| CP-d(GCG) | 3726120 | 2615084 | 1660563 | Sensitivity decreased 55.4% |
| CP-d(GTG) | 4003424 | 3739408 | 2342036 | Sensitivity decreased 41.5% |

**Table S3**: A 1 pmol aliquot of each CP-d(GpX) standard and IS was analyzed by the appropriate UPLC-SIM assay using 15 mM ammonium acetate at pH 7.0, pH 5.5, and pH 4.0 respectively. Overall change in sensitivity was calculated using the third-most abundant isotope (quantitative peak).

| Substrate | Ratio (Std:IS) | Accuracy | Precision | Linearity | LOQ | LOD |
|---|---|---|---|---|---|---|
| CP-d(AG) | 0.15:2 | 101.9% | 0.9% | $y = 1.0493x – 0.0048$<br>$r^2 = 0.9961$ | 50 fmol | 25 fmol |
| | 0.06:2 | 98.7% | 1.7% | | | |
| | 0.03:2 | 70.8% | 30.2% | | | |
| | 0.015:2 | 99.9% | 12.2% | | | |
| CP-d(GG) | 0.20:1 | 93.4% | 8.3% | $y = 1.0767x – 0.003$<br>$r^2 = 0.9984$ | 20 fmol | 10 fmol |
| | 0.08:1 | 103.8% | 5.8% | | | |
| | 0.05:1 | 94.6% | 24.1% | | | |
| | 0.025:1 | 108.0% | 5.6% | | | |
| | 0.01 | 105.0% | 40.0% | | | |
| CP-d(GCG) | 0.5:1 | 100.3% | 0.017% | $y = 1.0767x - 0.003$<br>$r^2 = 0.9984$ | 20 fmol | 10 fmol |
| | 0.25:1 | 106.3% | 3.2% | | | |
| | 0.1:1 | 91.7% | 11.1% | | | |
| | 0.05:1 | 82.1% | 4.0% | | | |
| | 0.02:1 | 75% | 9.4% | | | |
| CP-d(GTG) | 0.5:1 | 91.3% | 9.2% | $y = 0.916x + 0.0059$<br>$r^2 = 0.9955$ | 10 fmol | 5 fmol |
| | 0.25:1 | 101.7% | 9.8% | | | |
| | 0.1:1 | 86.0% | 7.2% | | | |
| | 0.05:1 | 109.8% | 16.7% | | | |
| | 0.02:1 | 103.7% | 19.8% | | | |

**Table S4**: Untreated CTDNA was spiked with a known amount of authentic standard and IS, followed by digestion by *Method 1* (CP-d(GG) and CP-d(AG)) or *Method 3* (CP-d(GCG) and CP-d(GTG)). After digestion, the enzymes were removed by Nanosep 10kDa filter centrifugation, CP-d(GpX) were enriched by offline HPLC, and then analyzed by the appropriate UPLC-SIM assay. The observed analyte:IS ratio was plotted against the theoretical ratio to establish linearity of assay. Accuracy of each point was calculated by subtracting the mean ratio by the expected ratio, followed by dividing by the expected ratio. The precision of each point was calculated by dividing the standard deviation by the mean ratio. The limit of detection of each assay was determined as a signal to noise ratio of 3:1, and the limit of quantitation was determined as a signal to noise ratio of 10:1 from the validation results.

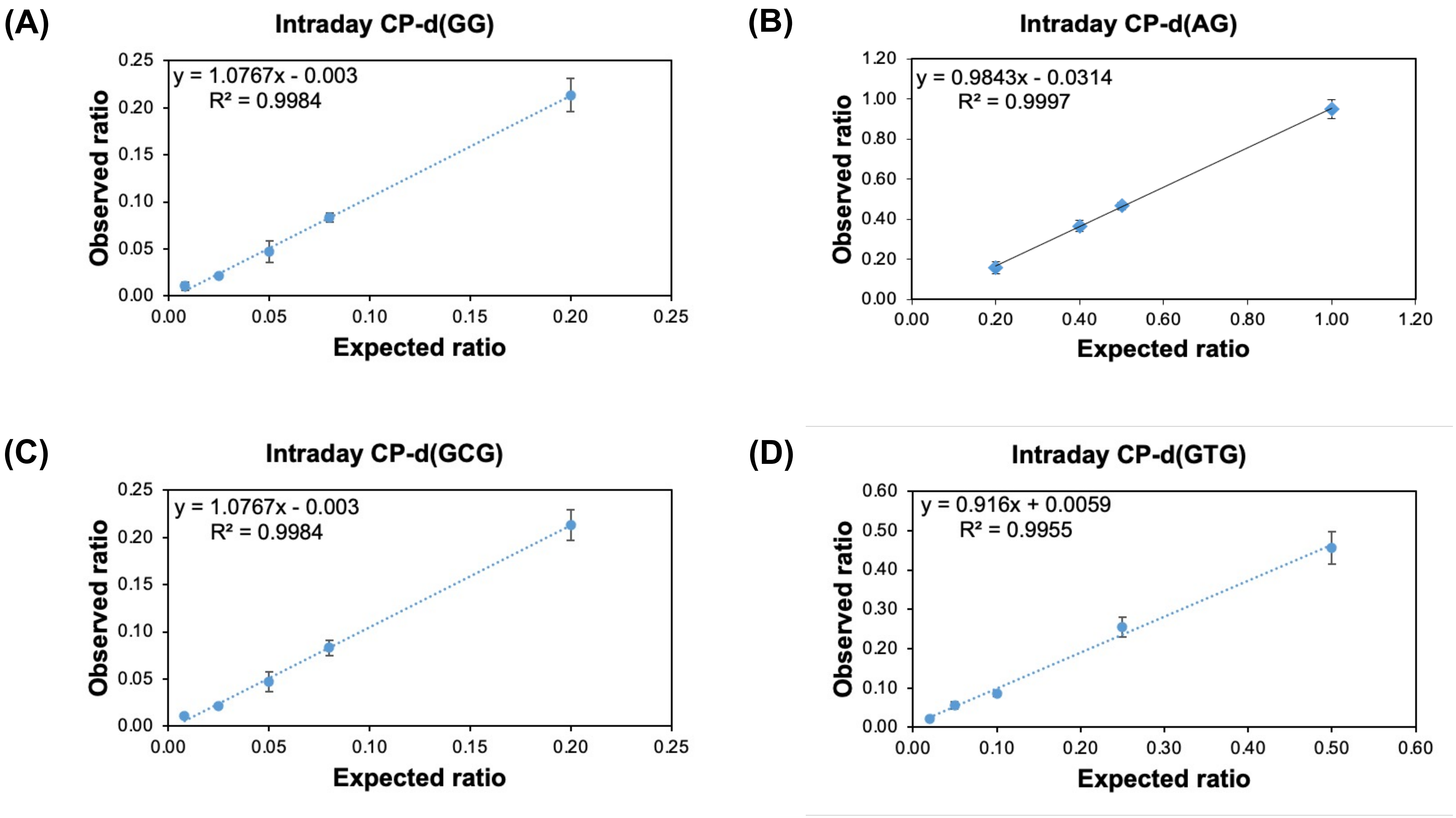


**Figure S4**: Graphs of the intraday validation results are shown for **(A)** CP-d(GG), **(B)** CP-d(AG), **(C)** CP-d(GCG), and **(D)** CP-d(GTG).

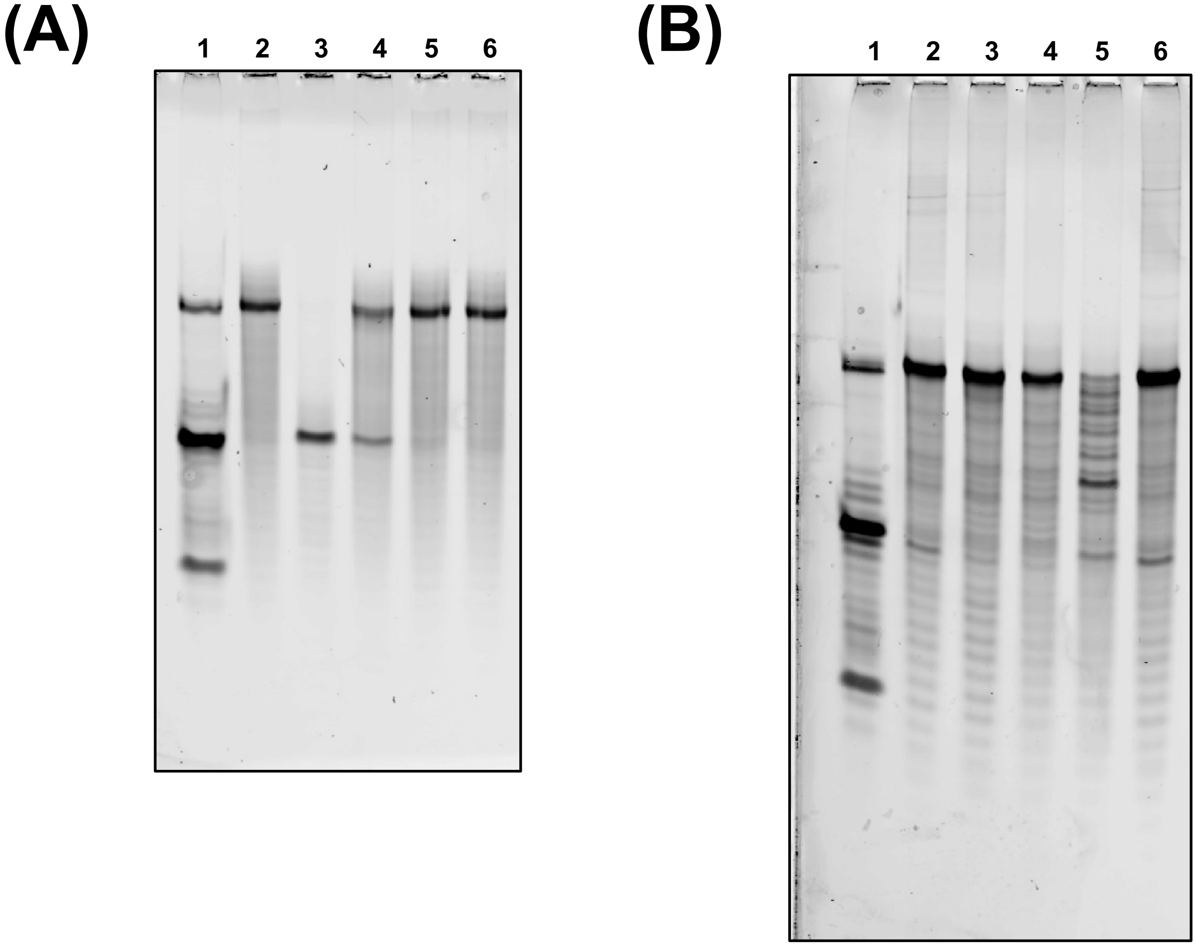


**Figure S5**: Optimizing digestion efficiency to yield CP-d(GpX). (**A**) Representative 20% urea gel of a 42mer 1,3-GTG-XL oligo digestion by Exo **I** to identify conditions where the enzyme can digest DNA up-to the cisplatin cross-link. **Lane 1**: DNA ladder of 10 pmol 42mer, 20mer, and 13mer. **Lane 2**: 10pmol 42mer 1,3-GTG-XL oligo. **Lane 3**: 42mer 1,3-GTG-XL oligo digested by 2 U ExoI. **Lane 4**: 42mer 1,3-GTG-XL oligo digested by 200 mU ExoI. **Lane 5**: 1,3-GTG-XL oligo digested by 20 mU ExoI. **Lane 6**: 1,3-GTG-XL oligo digested by 2 mU ExoI. **(B)** Representative 20% urea gel of a 42mer 1,3-GTG-XL oligo digested by Exo **III** to identify conditions where the enzyme can digest DNA up-to the cisplatin cross-link. **Lane 1**: DNA ladder of 10 pmol 42mer, 20mer, and 13mer. **Lane 2**: 42mer 1,3-GTG-XL oligo digested by 2 mU ExoIII. **Lane 3**: 42mer 1,3-GTG-XL oligo digested by 20 mU ExoIII. **Lane 4**: 42mer 1,3-GTG-XL oligo digested by 200 mU ExoIII. **Lane 5**: 1,3-GTG-XL oligo digested by 2 U ExoI. **Lane 6**: 10pmol 42mer 1,3-GTG-XL oligo.

| Enzyme | 1,2-GG-Intra XL | 1,2-AG-Intra XL | 1,3-GCG-Intra XL | 1,3-GTG-Intra XL |
|---|---|---|---|---|
| PDE II (5' – 3') | Full | Full | Full | Full |
| Exo I (3' – 5') | Partial | Full | Full | Full |
| Exo III (3' – 5') | Partial | Partial | Partial | Partial |
| PDE I (3' – 5') | None | None | None | None |
| Exo V (Both) | None | None | None | None |
| Exo VII (Both) | None | None | None | None |
| T7 Exo (5' – 3') | None | None | None | None |
| MNase (endo) | None | None | None | None |

**Table S5**: Table summarizing the results of 42mer oligo digestion by a single enzyme. The term "full" represents digestion up to the cisplatin cross-link, "partial" represents incomplete digestion yielding multiple products including digestion up the cisplatin cross-link, and "none" represents no desired product observed.

(A)

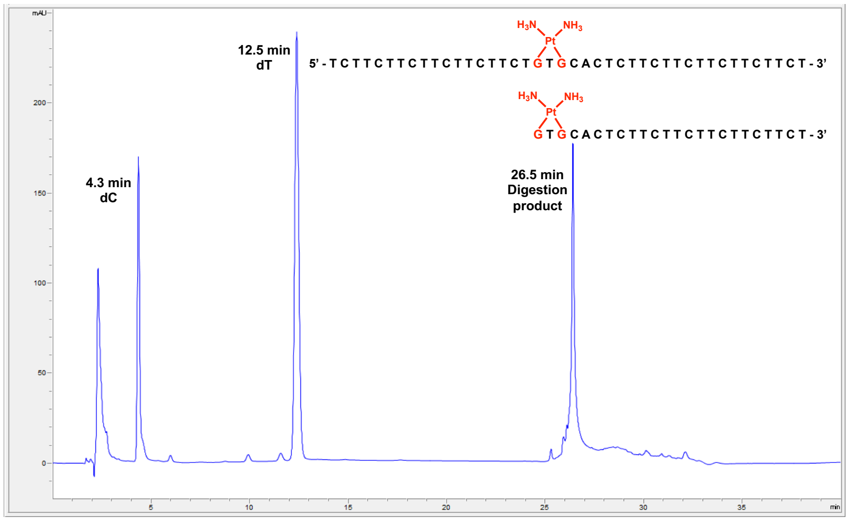


(B)

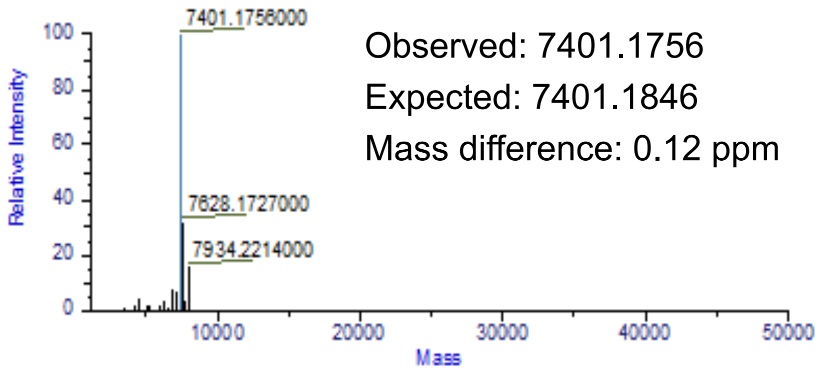


(C)

| Substrate | Enzyme | Observed mass | Expected mass | Mass difference |
|---|---|---|---|---|
| 42mer 1,2-GG Intrastrand XL | PDE II (5' – 3') | 7407.1729 | 7407.1755 | 0.35 ppm |
| | Exo I (3' – 5') | NA | NA | NA |
| | Exo III (3' – 5') | 6205.9899 | 6205.9807 | 1.48 ppm |
| 42mer 1,2-GTG Intrastrand XL | PDE II (5' – 3') | 7401.1756 | 7401.1846 | 0.12 ppm |
| | Exo I (3' – 5') | 6510.0359 | 6510.0348 | 0.17 ppm |
| | Exo III (3' – 5') | NA | NA | NA |

**Figure S6**: Detailed analysis of a single enzyme digestion of a 42mer cross-link substrate (**A**) Representative HPLC purification trace of 42mer 1,3-intrastrand cross-link oligo digestion product after incubation with 0.1 U PDE II. (**B**) Representative LC-MS confirmation of the 42mer 1,3-GTG-Intrastrand cross-link digestion product after incubation with 0.1 U PDE II. (**C**) Table summarizing the results of 42mer intrastrand cross-link digestion by a single exonuclease.

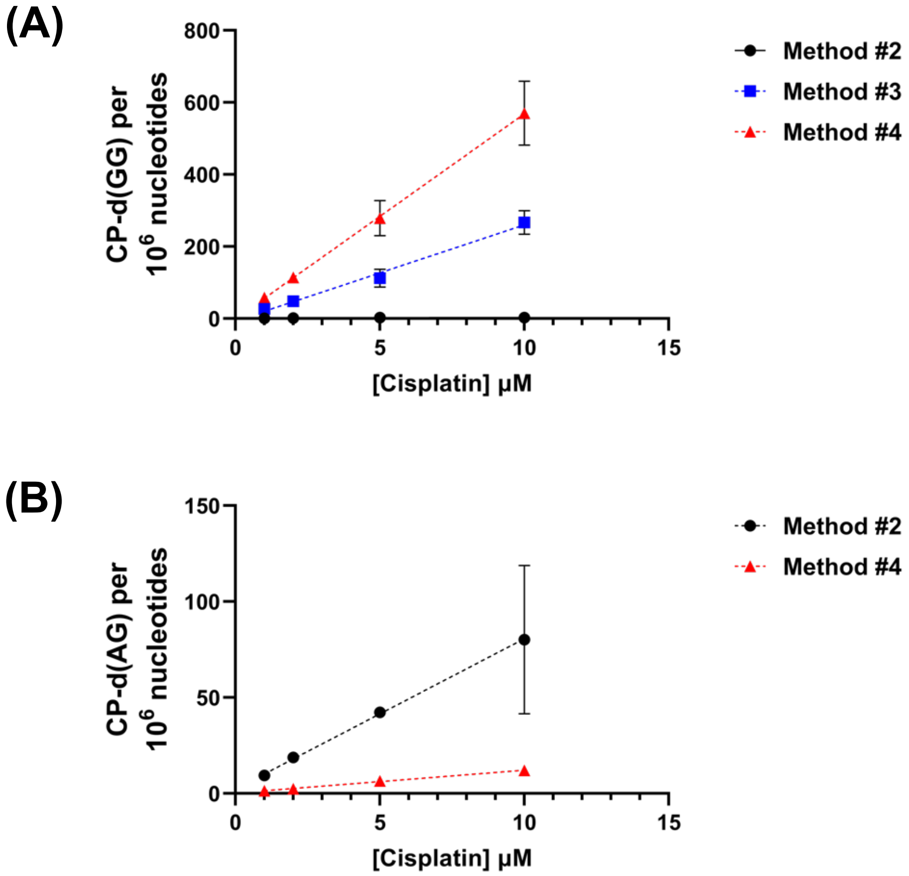


**Figure S7**: Evaluating alternative digestion methods of 1,2-intrastrand cross-links. **(A)** Comparison of *Methods 2 – 4* to quantify CP-d(GG) from CP-treated CTDNA. **(B)** Comparison of *Methods 2* and *4* to quantify CP-d(AG) from CP-treated CTDNA.

(A)

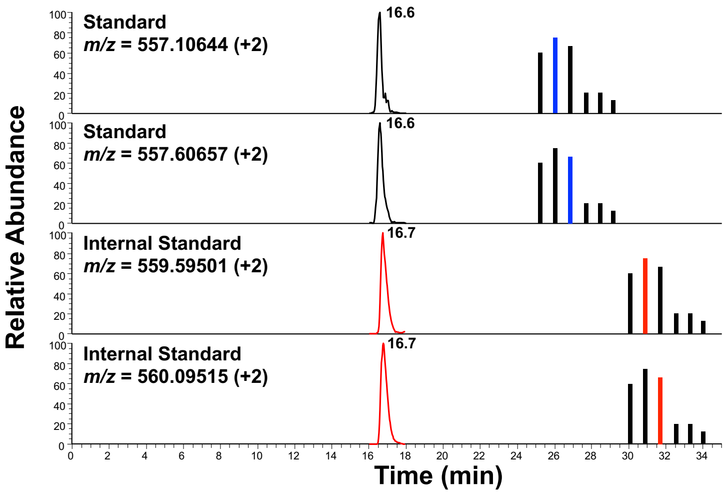


(B)

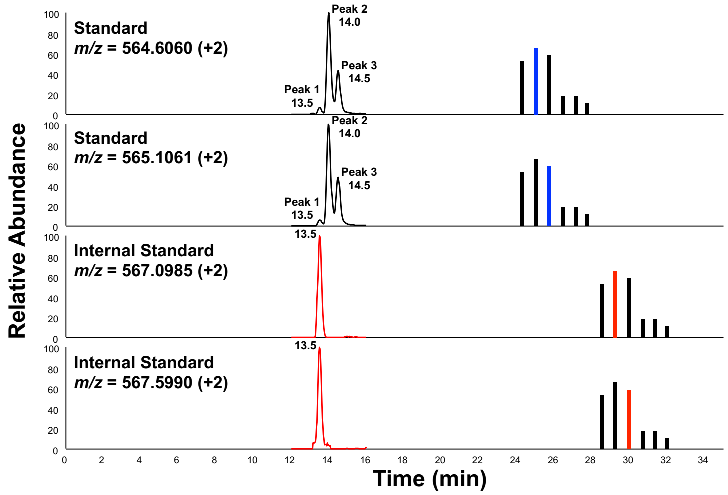


(C)

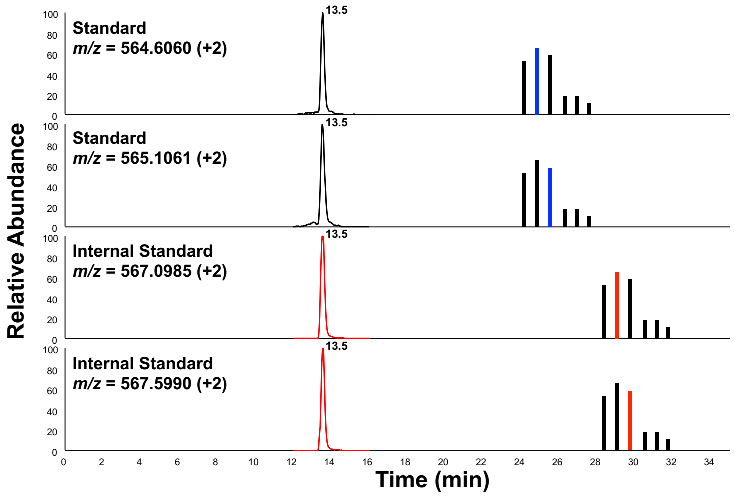


**Figure S8**: Representative traces of CP-d(GXG) obtained from Method #3 digestion. **(A)** CP-d(GCG) from 10 µM CP-treated CTDNA. **(B)** CP-d(GTG) from 10 µM CP-treated CTDNA. **(C)** Representative trace of CP-d(GTG) from CP-treated double-stranded 42mer oligo containing a single, internal GTG sequence (1:1 dsDNA:CP).

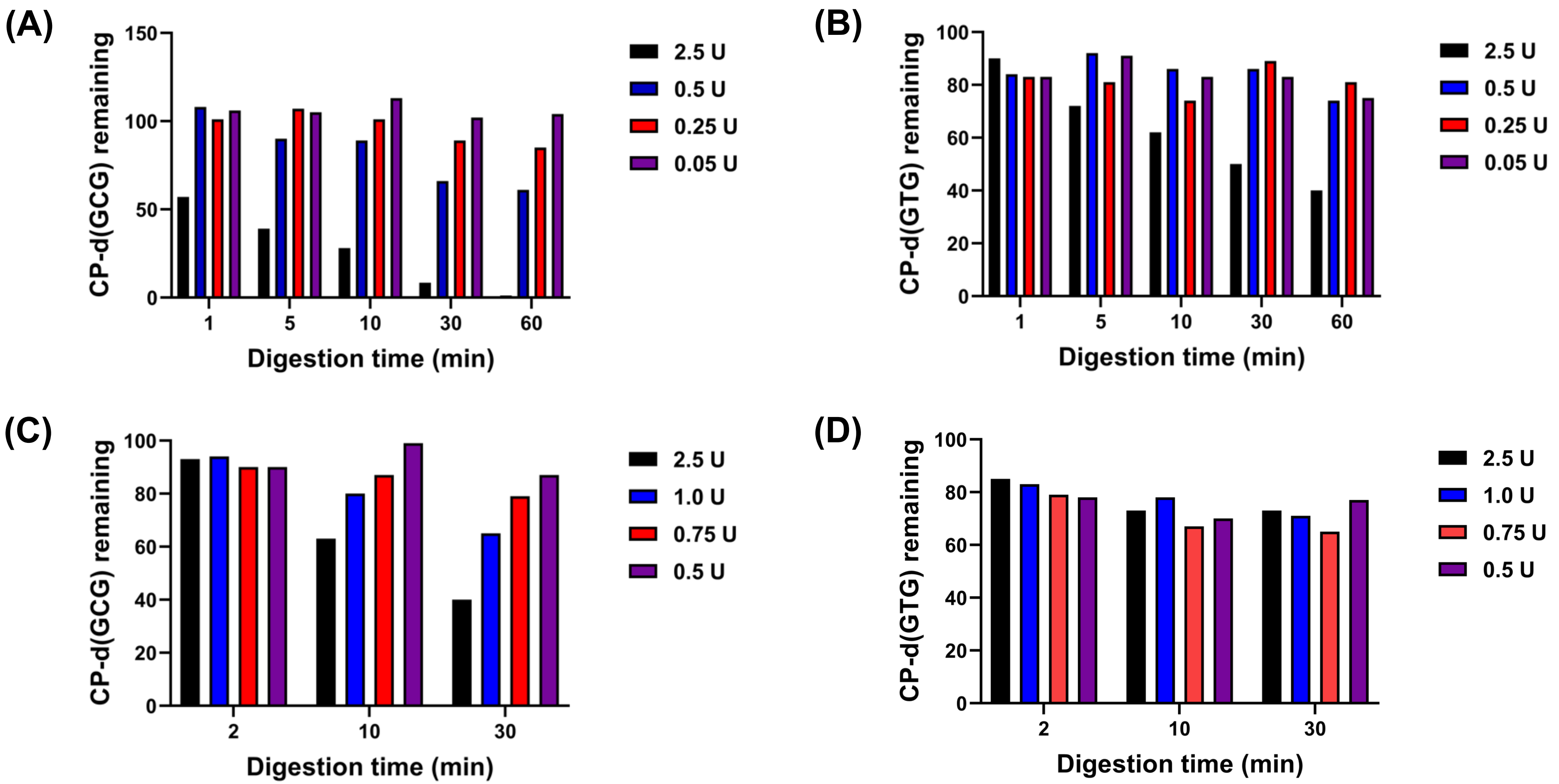


**Figure S9**: Optimizing *Method 3* digestion conditions to prevent over-digestion of CP-d(GXG). **(A)** Digestion results of CP-d(GCG) standard and IS spiked in reaction buffer and incubated with NEB nucleoside digestion mix only. **(B)** Digestion results of CP-d(GTG) standard and IS spiked in reaction buffer and incubated with NEB nucleoside digestion mix only. **(C)** Digestion results of CP-d(GCG) standard and IS spiked in 25 µg CTDNA and incubated with NEB nucleoside digestion. **(D)** Digestion results of CP-d(GTG) standard and IS spiked in 25 µg CTDNA and incubated with NEB nucleoside digestion.
